# Dynamic coexistence driven by physiological transitions in microbial communities

**DOI:** 10.1101/2024.01.10.575059

**Authors:** Avaneesh V. Narla, Terence Hwa, Arvind Murugan

**Affiliations:** Department of Physics, University of California, San Diego; Department of Physics, University of Chicago

## Abstract

Microbial ecosystems are commonly modeled by fixed interactions between species in steady exponential growth states. However, microbes often modify their environments so strongly that they are forced out of the exponential state into stressed or non-growing states. Such dynamics are typical of ecological succession in nature and serial-dilution cycles in the laboratory. Here, we introduce a phenomenological model, the Community State model, to gain insight into the dynamic coexistence of microbes due to changes in their physiological states. Our model bypasses specific interactions (e.g., nutrient starvation, stress, aggregation) that lead to different combinations of physiological states, referred to collectively as “community states”, and modeled by specifying the growth preference of each species along a global ecological coordinate, taken here to be the total community biomass density. We identify three key features of such dynamical communities that contrast starkly with steady-state communities: increased tolerance of community diversity to fast growth rates of species dominating different community states, enhanced community stability through staggered dominance of different species in different community states, and increased requirement on growth dominance for the inclusion of late-growing species. These features, derived explicitly for simplified models, are proposed here to be principles aiding the understanding of complex dynamical communities. Our model shifts the focus of ecosystem dynamics from bottom-up studies based on idealized inter-species interaction to top-down studies based on accessible macroscopic observables such as growth rates and total biomass density, enabling quantitative examination of community-wide characteristics.

## Introduction

Microbial communities in natural environments are often highly dynamic [1–7]. For example, many environments feature periodic replenishment of resources (e.g., the gut microbiome [8], the ocean [9]), or resetting of other environmental factors with periods of growth between these perturbations [10, 11]. Lab-scale experiments [12–14] on microbial ecosystems frequently adopt serial dilution cycles with dynamic environments. Recent studies have found that stable microbial communities do not settle simply into a fixed state, but are instead driven through dynamic phases involving complex changes in the environment such as depletion of oxygen and build-up of toxic waste [15, 16]. These changes, in turn, alter the physiological states of the microbes in the community, slowing down or even halting their growth. Changing physiological states also often change metabolic secretion and uptake profiles, and induce more complex interactions such as aggregation, motility, toxin secretion, and even contact-dependent killing [17–20].

These observations have been interpreted using models from theoretical ecology that typically explain ecosystem assembly and stability [21, 22] in terms of resource competition [12, 13, 23], niche differentiation [24] and competitive exclusion [25]. However, these models typically assume that communities and the organisms in them are at steady state [21, 26–29]. This difference between empirical observations and theoretical models raises questions about the role of dynamic physiological state changes in forming complex communities. One possibility is that physiological state changes are merely details, not essential for understanding factors that enable community assembly. In this perspective, nothing is lost by coarse-graining over dynamics and modeling communities as if they are at steady state. Microbial communities would be expected to show similar complexity and structures if microbes stay in fixed states (e.g., exponential growth) and independent of whether interactions through metabolic secretion and uptake occur in a temporally staged manner.

Another possibility is that physiological state changes create dynamic niches that support complex communities. Since microbes have a plethora of non-growing states, this scenario could significantly expand the ways of generating niches beyond well-studied cases such as distinct metabolites [28], space [30], and externally dictated temporal epochs (e.g., diel or annual cycles) [31]. Further, the nature of such self-generated dynamic niches, if they exist, might have signatures that are predicted to be observed in microbial communities.

We cannot easily address this question about the role of state changes using the current bottom-up theoretical frameworks (e.g., Lotka-Volterra or Consumer-Resource models) since these models typically characterize organisms and their interactions with fixed parameters. In these models, community dynamics only involves changes in species abundances and nutrient concentrations and is justified by assuming organisms are in a fixed physiological state, (e.g., Monod growth for exponentially-growing cells [32]). If one is to adopt a model of interactions between each species and its environment (as in Consumer-Resource Models) or other species (as in Lotka-Volterra Models), then each physiological state would minimally involve a different set of uptake and excretion parameters; a given species would effectively be modeled as multiple species over time. Thus, bottom-up models of dynamic communities require extensive characterization and unconstrained assumptions on specific details about what different cells do in different conditions.

As a first step towards quantitatively modeling communities of species that undergo physiological changes, we introduce a minimal top-down phenomenological framework, the Community State Model. Our model is phenomenological at the level of species density; the physiological state and thus the growth rate of each species in a community is assumed to depend only on the community biomass at any given time, and as a result, community states are defined by regions of biomass density. Such a model can be solved explicitly (numerically and in simple cases analytically) to yield the temporal organization of community dynamics at a quantitative level.

Analysis of the Community State model points to sequential dynamics as a strategy to form a stable community involving a large number of species [33]. In this simple model, each species grows rapidly in one (or a few) community states that persist over specific intervals of biomass accumulation, with slower or even no growth in other parts of the inter-dilution period (hereby referred to as the growth period). This strategy is a distinct alternative to the co-growth strategy based on steady-state models with fixed physiological states where species grow on resource niches simultaneously.

For this sequential coexistence strategy, our model allows us to uncover a number of key features of community dynamics. We find (a) tolerance of community diversity to fast-growing species if such growth is limited to specific community states, (b) enhanced community stability through staggered dominance of different species in different community states, and (c) a requirement of increased growth dominance for late-growing species. These features counteract the dominant notions regarding species competition derived from analysis of steady-state systems, and serve as principles to guide the understanding of complex dynamical ecosystems

## Results

### Case study of community dynamics during serial-dilution cycles

The model of microbial community dynamics developed here is inspired by dynamics revealed by a recent study of a seemingly simple cross-feeding system subjected to repeated serial-dilution cycles [16]: Two species of marine bacteria, *Vibrio sp. 1A01* and *Neptunomonas sp. 3B05*, isolated from a chitin-degrading coastal community [34], were grown on N-acetyl glucosamine (GlcNAc) as the sole carbon and nitrogen sources in batch culture. 1A01 consumes GlcNAc and excretes acetate and ammonium, while 3B05, which does not consume GlcNAc, can grow on the acetate and ammonium excreted by 1A01 (Fig. 1A). Because 1A01 grows on GlcNAc faster than 3B05 grows on acetate, acetate inevitably accumulates in the medium, becoming toxic when reaching the buffer capacity of the medium (which is low for seawater). In canonical syntrophy, toxicity would slow down the toxin-excreting species more than the toxin-clearing species, resulting in a stable state where both species grow exponentially [35]. This is not the case for 1A01-3B05 co-culture: acetate accumulation in the environment slows down the growth of 3B05, the acetate-consumer, more than that of 1A01, the acetate excretor. Thus acetate is expected to accumulate, with the arrest of the co-culture once acetate exceeds the buffer capacity. Yet, as shown in Fig. 1B, when this system was subjected to 24-h growth-dilution cycles, the system miraculously cured itself of acetate accumulation, with the two species reaching a stable, comparable abundance ratio according to samples taken at the end of each cycle after a few cycles.

**Figure 1:**
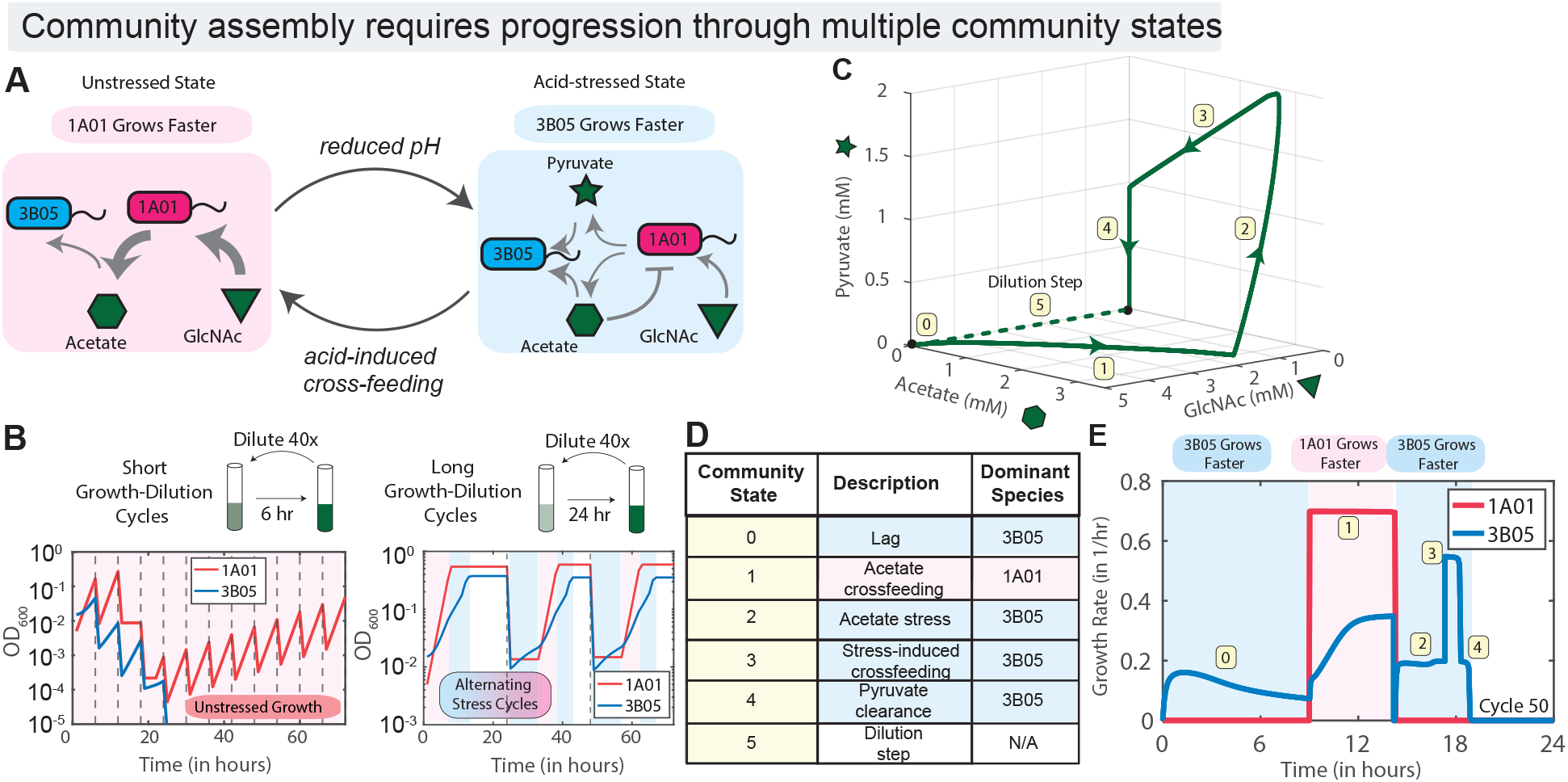
Sequential transitions in a community’s state revealed in a simple co-culture of a sugar consumer and an organic acid consumer. (A) Schematics describing the dynamics of a co-culture of marine bacteria *Vibrio sp. 1A01*, a sugar consumer, and *Neptumonas sp. 3B05*, an organic acid consumer, growing on N-acetyl glucosamine (GlcNAc) under repeated serial-dilution cycles: the co-culture passes from an initial unstressed state in which 1A01 grows faster than 3B05 (pink box), to an acid-stressed state in which 3B05 grows but 1A01 does not (blue box). Pointed gray arrows indicate metabolic flow (thickness indicates flow magnitude) and blunt-end arrows indicate growth inhibition. (B) Experimental investigation by Amarnath et al. [16] revealed coexistence only if serial-dilution cycles were sufficiently long to allow for an intricate sequence of ‘community states’, i.e., different combinations of the physiological states of each species and media conditions, labeled by the numbers in (C,D). (C) shows changes in 3 major metabolites while Table (D) describes the community states along the green path in (C). (E) The growth rate of each species along the path in (C), i.e., between two serial dilutions (steady state cycle shown).

Detailed analysis of the two-species dynamics revealed that stability was achieved through physiological transitions during the 24-h growth period, as shown in Fig. 1C-E: The first phase 🄋 after nutrient replenishment is a lag phase for 1A01; in the next phase ➀, acetate accumulated and pH dropped; 1A01 grew fast with slower growth for 3B05. Phase ➁ commenced when the pH hit a critical threshold (set by the pKa of acetic acid) which caused 1A01 and 3B05 to enter growth arrest, with 1A01 excreting large amounts of glycolytic intermediates (e.g., pyruvate). This stress-induced excretion played a key role in stimulating the growth of 3B05 (phase ➂ ), with the consequence of removing acetate from the medium and restoring pH (phase ➃). This in turn allows 1A01 to avoid death and be available for dilution into the next cycle (phase ➄ ).

Amarnath et al.[16] showed that this highly dynamical mode of coexistence is not specific to the marine species studied: dynamic coexistence through similar acid shock and recovery was shown also for co-culture of species taken from a soil community or even between enteric and soil bacterium. Metabolic analysis in [16, 36] suggests that such interactions are generic between species with complementary sugar-preferring vs. acid-preferring bacteria, or between glycolytically-oriented vs. gluconeogenically-oriented modes of metabolism. Thus, dynamic coexistence with each species passing through multiple physiological states in a cycle may be the norm rather than the exception [12, 37–39]. The lack of reports of such dynamical features may reflect the lack of time-resolved measurements, which occurred within a few-hour window of the 24-hour growth-dilution cycles. The focus of existing theoretical studies in ecology on steady-state characteristics and stable coexistence of many species [22, 40] may also contribute to the lack of measurements on dynamic characteristics.

### Benefits of indexing community dynamics by community biomass

How can we effectively capture the key drivers of the complex dynamics underlying this system? We propose a phenomenological and experimentally accessible description, using “community biomass” as a proxy for key drivers of community dynamics.

One advantage of this biomass-based description is that the community state changes at reproducible values of the accumulated biomass for perturbations in external parameters. This robustness arises because changes in environmental parameters like pH and oxygen levels are typically accompanied by biomass accumulation. In contrast, the most direct description based on time has several drawbacks. As shown in Fig. 2A, community dynamics during a growth cycle in real time are highly variable as initial species or nutrient abundances are varied. However, this variance is mostly counteracted by the variation in the real-time accumulation of biomass; Fig.2B. Hence, combining Fig. 2A and 2B, we find that the growth rate as a function of accumulated biomass is relatively reproducible; see Fig.2C. See *Supplementary Supplementary Figures S1-2* for other ecosystems with even higher reproducibility of biomass. While biomass values corresponding to community state changes will be different when, say, a given species is part of a novel community, these results suggest that total biomass could be part of a useful top-down description of a given ecosystem.

**Figure 2:**
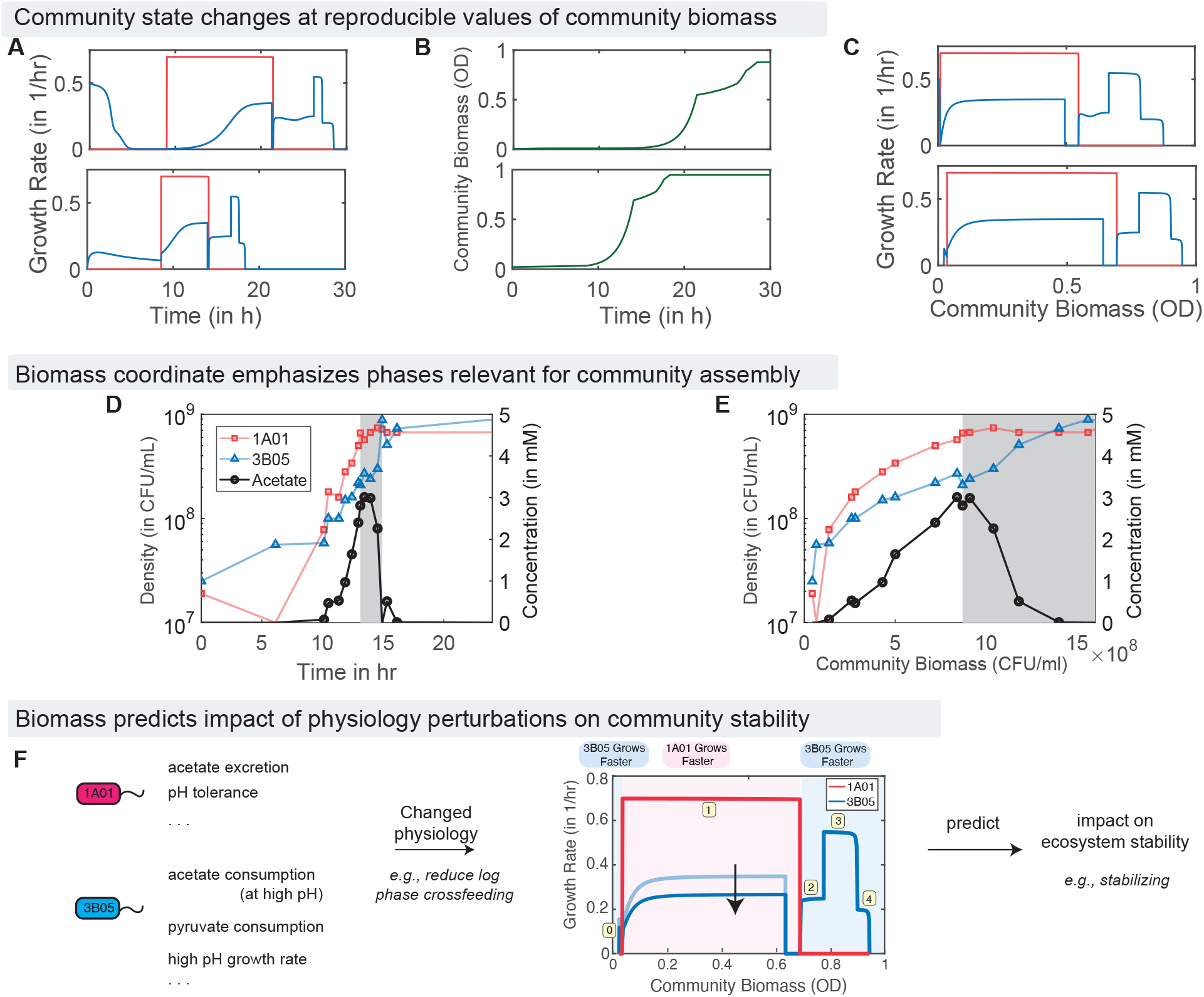
Benefits of indexing community states by the accumulated community biomass. When plotted as a function of time during a growth period, different initial conditions (here, relative species abundances of A:B=1 for top row, A:B=0.66 for bottom row) for the system in Fig. 1 lead to (A) different growth rate curves of each species and (B) accumulated community biomass density. (C) However, growth rate data as a function of community biomass density is relatively reproducible. Results shown from second cycle (before reaching stable cycle). (D,E) Experimental data from [16] on cell densities and acetate concentration during a growth dilution cycle. (D) The most important physiological changes that underpin coexistence in [16] occur in a short window of time (gray) where acid stress causes dramatic changes in acetate concentration (black curve) and other metabolites (not shown). However, without prior knowledge of the acetate stress mechanism, sampling uniformly in real-time will dedicate many time points to the lag phase (relatively unimportant for coexistence [16]) and may miss the critical gray region. (E) In contrast, sampling evenly in community biomass naturally emphasizes the gray region. Thus investigating metabolites that change dramatically between biomass density intervals can assist in identifying the mechanistic basis of community assembly. (F) A mutation that changes physiology in specific environments (here, reducing log phase crossfeeding) will change growth curves in community biomass intervals corresponding to those environments (here, the light blue growth curve for 3B05 is lowered to the dark blue curve in the pink biomass interval). In this work, we derive a formula relating growth rate curves as a function of biomass to coexistence. Consequently, our top-down framework can relate changes in physiological properties to community assembly.

Another advantage of indexing the community dynamics by total biomass is that it naturally emphasizes growth phases most relevant for community assembly. As shown in Fig. 2D, when plotted in real time, the most critical physiological changes occur in a relatively brief period indicated by the gray band (where acetate buildup hits a threshold, leading to subsequent stress-induced crossfeeding of pyruvates and other metabolites). There would have been no reason to sample the short time period represented by the gray band in Fig. 2D, without data on acetate (black curve) and other results of the detailed mechanistic study in [16]. More generally, the drawback of real time is because environmental change is a result of *absolute* biomass growth, and for an exponentially-growing culture, the same absolute amount of environmental change takes exponentially less time as the culture grows. Instead, sampling the ecosystem uniformly in accumulated community biomass density (Fig. 2E) emphasizes important periods such as the gray region, even if the role of acid stress was unknown.

Finally, as we will detail later, the accumulation of sufficient biomass is the necessary and sufficient requirement for each species to be maintained in stable cycles. Consequently, we can use this picture based on biomass to predict the impact of, e.g., altered physiology in an organism due to mutations, on coexistence. For example, consider a change that decreases cross-feeding during exponential growth as shown in Fig. 2F. Naively, such a change would be expected to destabilize coexistence as cross-feeding is believed to enhance coexistence [12]. But our biomass-based results derived below will predict the opposite; the depicted physiological change causes 3B05 will grow slower during 1A01’s growth phase. We will show that reducing co-growth in intervals of the biomass coordinate will generally favor coexistence since each species will get guaranteed (but capped) growth in the community state that it dominates. Consequently, our framework will predict reducing cross-feeding during the exponential phase can stabilize the community, counter to intuition.

### A top-down model for complex communities

We propose a general model - the Community State (CS) Model - for investigating multi-species dynamics in microbial communities in cyclic environments. In this top-down model, we take the growth rate, *r*_*α*_(*S*), of each species *α* to depend on the community state *S*, which progresses through multiple states due to various environmental changes driven by the microbes themselves as shown in Fig. 3. In the simplest model, we assume that the community state *S* can be parameterized by the community biomass, taken to be the total cell density 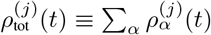, where 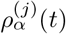 is the cell density of species *α* at time *t* in the *j*^th^ cycle. (Here, we assume all species to have the same biomass per cell. More generally, a scaling factor can be introduced to absorb the species-dependent cell mass.) Thus, the growth of species during the *j*^th^ cycle is described by

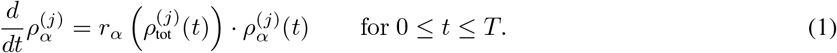

**Figure 3:**
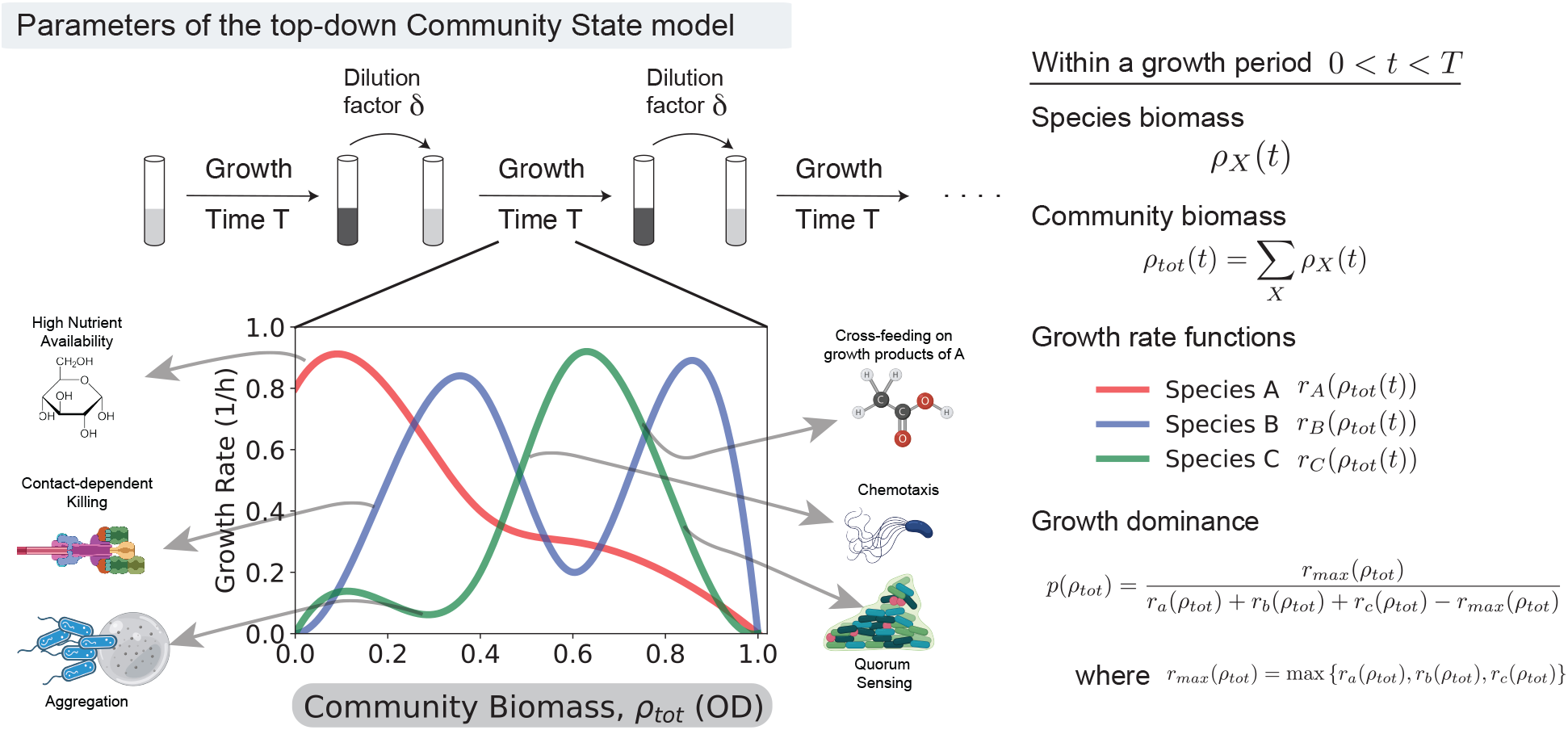
Top-down description of ecosystems in the Community State Model. We consider ecosystems that grow, gaining biomass for a period *T*, before being diluted into a fresh environment. During the growth period, the community changes its environment through many complex processes shown, leading to a sequence of community states. The Community State model assumes these processes to be turned on and off at different points along a one-dimensional phenomenological coordinate that parameterizes the sequence of community states. In the simplest version of the model shown, this coordinate is taken to be the accumulated community biomass *ρ*_*tot*_(*t*) since nutrient depletion, buildup of toxins, and spatial structure, etc. are accompanied by biomass growth. Each species is assigned a different set of growth rates *r*_*X*_ (*ρ*_*tot*_). At any given stage of the growth cycle, indexed by *ρ*_*tot*_, growth dominance *p*(*ρ*_*tot*_) is the ratio of the growth rate of the fastest-growing species to that of other species. As suggested by the cartoon, different species might dominate at different *ρ*_*tot*_ and to different extents *p*(*ρ*_*tot*_).

When *t* reaches the growth period *T*, all densities are reduced by a common factor *δ <* 1, i.e.,

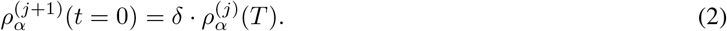

Eqs. [1] and [2] define an effective “map” for the density of each species at the beginning of each cycle, 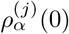 starting from the initial composition 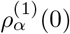. For convenience, we choose the growth period *T* to be sufficiently long such that the total cell density has time to reach the maximum 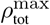 where *r*_*α*_ = 0 for all species. Thus, the dynamics of this model are governed by the growth rate functions *r*_*α*_(*ρ*_tot_), the maximal cell density of the system 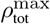, and the dilution factor *δ <* 1, independent of the growth period *T* .

The assumption that growth rates depend only on accumulated community biomass can be justified mechanistically in some cases; see *Supplementary Text S2*.*2 and Supplementary Figures S1-2*. But from a phenomenological point of view, this assumption can be viewed as the minimal closure of the equations describing an ecosystem that results in a coexistence criterion. Such a coexistence criterion is derived in *Supplementary Text S2*.*6* for mutual invasibility between two species. E.g., for the ability of one species, *A*, to invade a monoculture of species *B*, we find

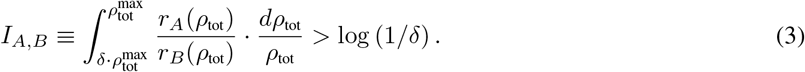

Similarly, the invasion of a species *A* monoculture by species *B* requires *I*_*B,A*_ *>* log (1*/δ*).

Intuitively, these conditions suggest that coexistence imposes conditions on the relative growth rate *r*_*A*_(*ρ*_*tot*_)*/r*_*B*_(*ρ*_*tot*_) in different intervals of community biomass *ρ*_*tot*_. For example, consider the growth curves *r*_*A*_(*ρ*_*tot*_), *r*_*B*_(*ρ*_*tot*_) shown in Fig. 4A. Species *A* grows faster than *B* for low biomass *ρ*_*tot*_ *< ρ*_*c*_, with a growth dominance *p*_1_ = *r*_*A*,1_*/r*_*B*,1_. Conversely, species *B* grows faster for higher biomass *ρ*_*tot*_ *> ρ*_*c*_, with growth dominance *p*_2_ = *r*_*B*,2_*/r*_*A*,2_. Applying Eq. **3** to this toy model with step-like growth curves as shown in Fig. 4A and C, we find conditions for mutual invasibility and thus for stable coexistence:

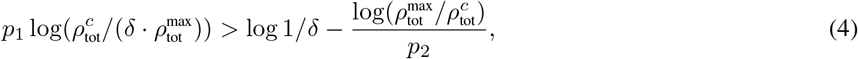

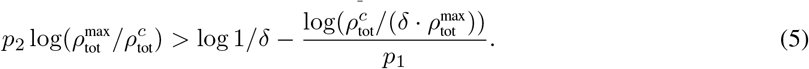

**Figure 4:**
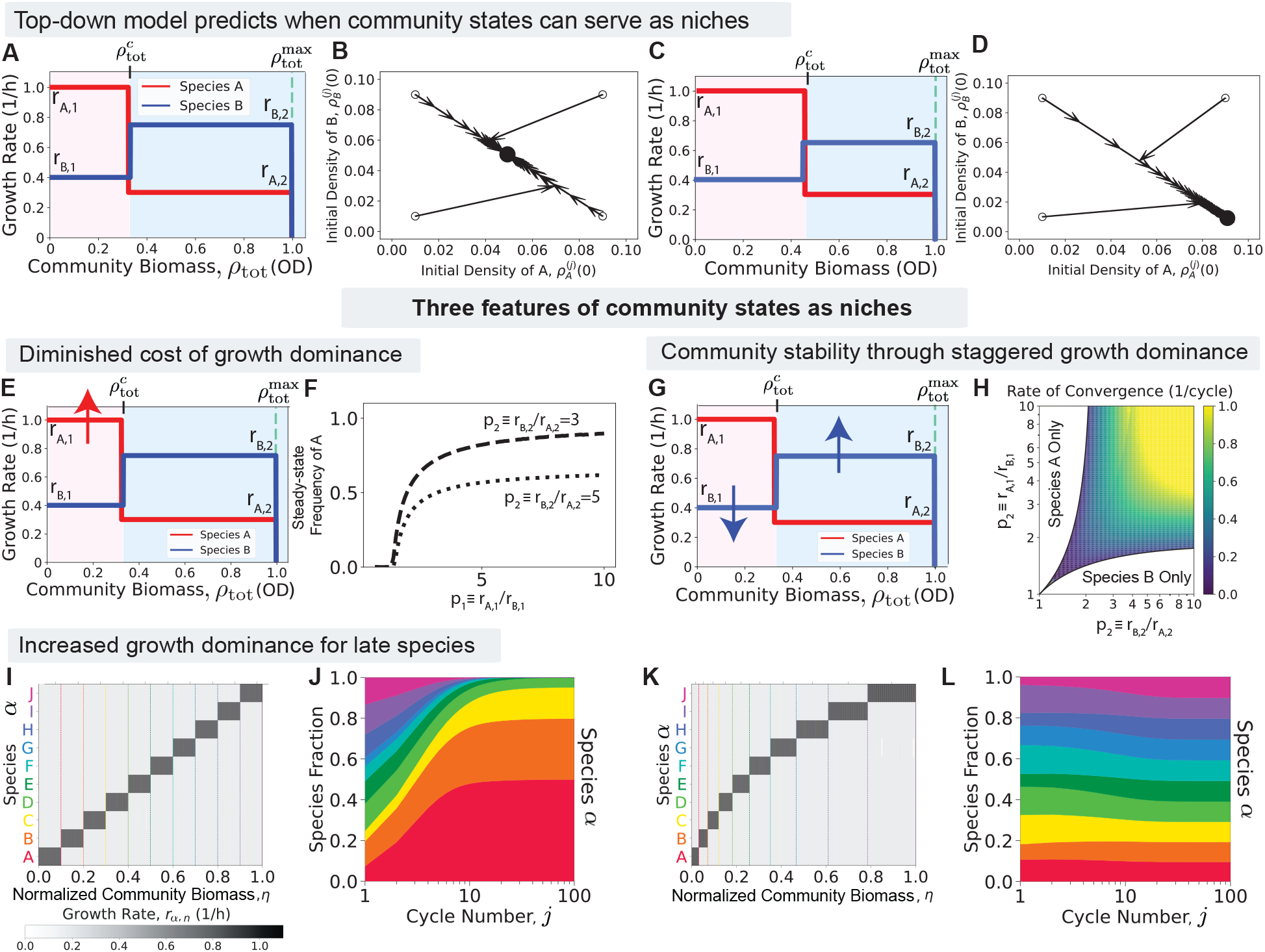
Key features of community states as niches. (A) Growth curves *r*_*X*_ (*ρ*_*tot*_) for two species; species *A* grows rapidly in the early community state that lasts for community biomass *ρ*_*tot*_ *< ρ*_*c*_; species *B* grows faster when *ρ*_*tot*_ *> ρ*_*c*_. (B) Species densities at the end of consecutive serial dilutions from different initial conditions (black open circles) for growth curves in (A) converge to a stable coexistence point (black solid circle). (C,D) Same as (A,B) but the modified growth curves in (C) lead to extinction of species *A* shown in (D). While growth curves in (A),(C) are visually similar, Eq.4,5 predict that the two community states can support two species in (A) but not in (C). **Key features:** (E) We increase growth dominance *p*_1_ = *r*_*A*,1_*/r*_*B*,1_ of species *A* in the first community state. (F) Steady-state abundance of species *A* as a function of growth dominance *p*_1_ shows a quickly saturating effect. (G) We consider a coordinated change of growth dominance *p*_1_, *p*_2_ across multiple community states. (H) Rate of convergence back to stable coexistence point (i.e., Lyapunov exponent of stability) as a function of growth dominances *p*_1_, *p*_2_; ecosystem stability increases without saturation for coordinated changes in dominance. (I) A 10-species ecosystem with each species “dominant” with high growth rate *r*_+_ in a unique community state and *r* elsewhere; all community states last equal intervals of normalized community biomass 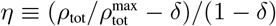. (Here *δ* is the dilution factor between growth cycles. Growth dominance *p* = *r*_+_*/r*_*−*_ = 5.) (J) Stacked chart shows species abundance at the end of each cycle (in fraction of total biomass) over multiple serial-dilution cycles. (K,L) Same as (I,J) but with wider biomass intervals for later community states (see Eq.6). The wider intervals for later species compensate for the priority effect enjoyed by early species.

When the growth dominances *p*_1_, *p*_2_ and the width of biomass interval 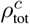 satisfy these conditions (Eq. **4**, Eq. **5**), serial dilution cycles starting from any initial condition converge to a unique fixed point where both species are present; see Fig. 4B. Outside of this regime, e.g., with visually similar growth curves shown in Fig. 4C, one of the two species goes extinct over multiple cycles as shown in Fig. 4D. See *Supplementary Text S2*.*4* for more analyses.

It is tempting to interpret community states in the two intervals of biomass shown in Fig. 4A as two dynamic ‘niches’ where species *A* and *B* dominate respectively, thus guaranteeing their coexistence. However, as the distinct steady-state compositions for similar growth curves in Fig. 4A and C show (coexistence and competitive exclusion respectively), community states can serve as distinct niches that support distinct species only under specific conditions. Thus the nature of niches that arise from community states remains unclear – what is the impact of the width of biomass intervals over which a community state persists, what effect does their temporal ordering have, and can community states support complex communities with many species? The phenomenological Community State model can be used to address these questions and makes non-trivial predictions on how growth and non-growth states must be structured across a community for stable coexistence. Below, we highlight three insights derived from these equations into the nature of niches based on community states:

#### Tolerance to growth dominance

As shown in Fig. 4E and F, the growth dominance *p*_1_ of species *A* in the first biomass interval negatively impacts the steady state coexistence ratio at small *p*_1_ but saturates at larger values, being limited by the value of the other growth dominance, *p*_2_. This diminishing damage to coexistence by a species with strong growth dominance can also be quantified by the tolerance range of parameters like *ρ*_*c*_ that allows for coexistence (see Supplementary Figure S3), follows from the structure of Eq. **3**.

The tolerance to growth dominance contrasts sharply with one of the most basic tenets of ecology, that faster-growing species drive slower ones to extinction, which is at the root of the “paradox of the plankton” [28]. It allows individual species with significant growth advantages to coexist with other slower species (thus retaining “services” by the latter in other more challenging community states), provided that these advantages are limited to some specific physiological states. This effect plays a pivotal role in the coexistence of larger communities to be described below.

#### Community stability through staggered growth dominance

While coexistence tolerates strong individual growth dominances as described above, the coexistence regime is broadened if *both p*_1_ and *p*_2_ are large, i.e., if the species stagger their dominance in distinct community states. The impact of staggered dominance is seen also in the robustness of coexistence, measured by the convergence rate to the stable cycle following small perturbations (the Lyapunov exponent); see Fig. 4G, H. This metric is also a measure of the stability of the ecosystem against environmental or physiological fluctuations.

Since enhancement in the size and stability of the coexistence region requires increases in growth dominance in *distinct* community states (i.e., *p*_1_ and *p*_2_), such a communal effect is aided by different species coordinating their growth dominance across multiple community state. However, individual species can also contribute to such staggered growth dominance since a species can increase the growth dominance of another species in another community state by reducing its own growth rate in that state; e.g., species *A* can increase *p*_2_, the dominance of species *B* in state 2, by decreasing *r*_*A*,2_. Thus, this global coordination is facilitated by individual species *specialized* to dominate in distinct community states.

#### Increased growth dominance for late species

The coexistence criteria *I*_*A,B*_, *I*_*B,A*_ are not symmetric between species *A* and *B*. This asymmetry can be traced to a “priority effect” where species *A* capitalizes on early growth, accumulating large numbers while species *B*’s growth occurs in a later biomass interval [41–43]. The consequences of this priority effect are shown for a *N* -species community in Fig. 4I,J: if each species is dominant in a unique biomass interval of equal width (with growth dominance *p* = 5 in each community state), late-growing species are driven extinct after several cycles.

We find that members of such a community can all coexist despite the priority effect if early-growing species occupy narrower biomass intervals. Based on Eq. **4** and Eq. **5**, we were able to derive a special distribution of biomass interval (see *Supplementary Text S2*.*8*) for coexistence. Expressed in the normalized biomass coordinate *η* = *ρ*_*tot*_*/ρ*_*max*_, if the width interval Δ*η*_*n*_ for the species growing in the *n*^th^-state follows an exponential distribution

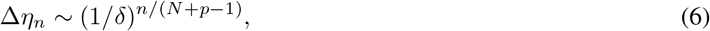

then late species coexist with early species and are, in fact, equi-abundant (provided that growth dominance *p* does not greatly exceed *N* ); see Fig. 4K, L. Thus, an appropriate choice of transition points between community states favoring the late-growing species can counteract the priority effect. Alternatively, the priority effect can be counteracted by stronger growth dominance of the late-growing species even if the biomass intervals of the different states are equal; see Supplementary Figure S6. These results underscore and quantify the large (exponentially increasing) burden faced by the late-growing species to be included stably in large communities.

### Complex communities

We next consider a multispecies Community State model in which different species can grow (and hence compete) in the same community state. This model can be represented by a “preference matrix” where each element of the matrix denotes the community states in which each species grows preferentially and the boundaries of the community states are common to all species and given by defined biomass values (*η*_*i−*1_ and *η*_*i*_ for the ith community state). We allow for multiple species to grow preferentially in any community state. This structure allows us to see the effect of competition between species growing preferentially in the same community states.

We first explored the case where each community state can support the rapid growth of a fixed number *K*_*s*_ of species. An example of a preference matrix is shown in Fig. 5A; each community state was assumed to last for the same average interval Δ*η* of biomass. In each community state (i.e., column of matrix shown), we assigned a high growth rate *r*_+_ for *K*_*s*_ randomly chosen species and a low growth rate *r*_*−*_ that is a *p* = 100 times lower than *r*_+_ for all other species. 30% random variation added to each parameter; see Supplementary Information for other details. The results, shown in Fig. 5B), show an expected decrease in steady-state diversity with an increasing number of species *K*_*s*_ competing in each community state. However, the decrease in surviving species is slower than a naive expectation of *N/K*_*s*_.

**Figure 5:**
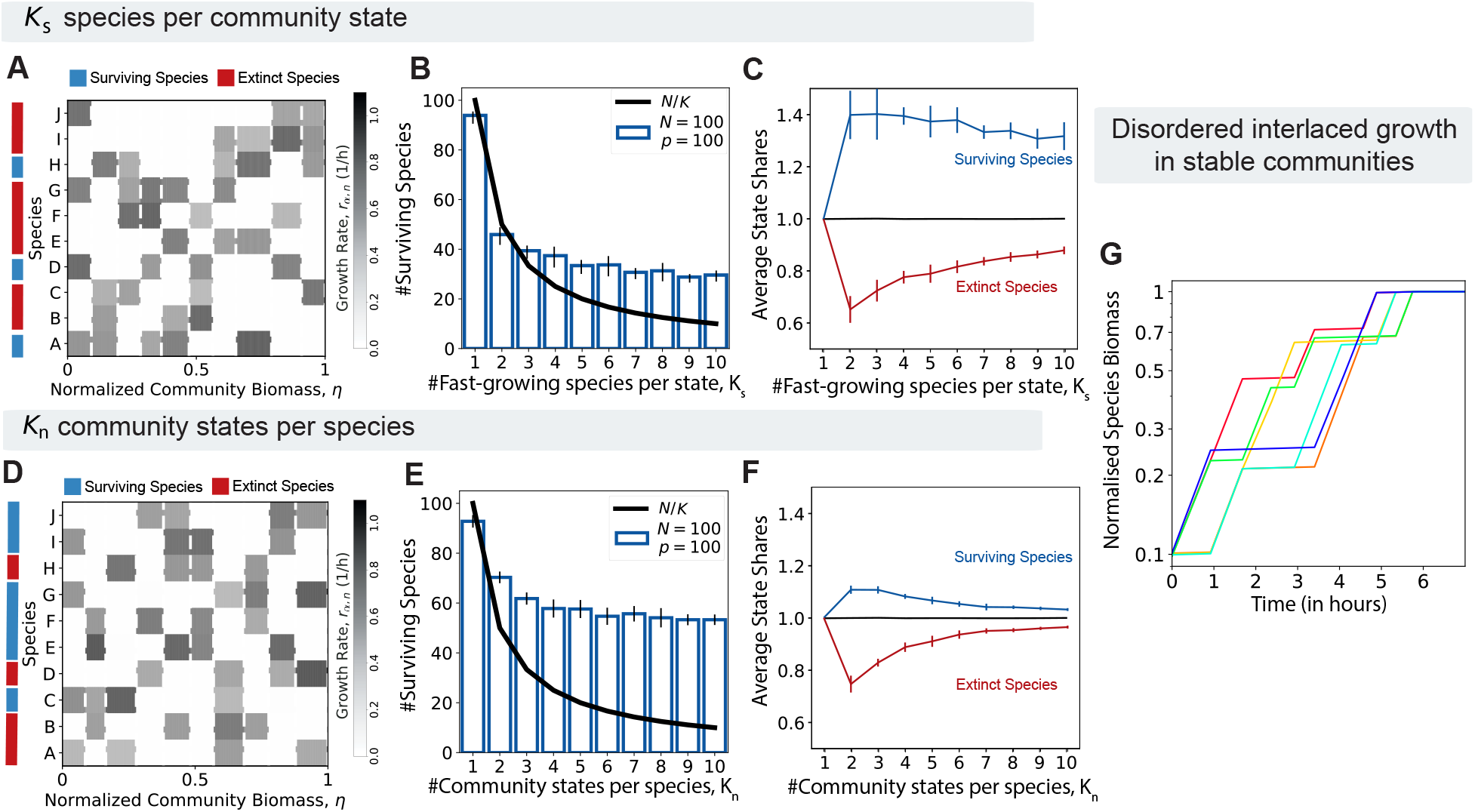
Competition and coexistence in complex communities. (A) *K*_*s*_ model of random preferences: *K*_*s*_ randomly chosen species are given high growth rate *r*_+_ in each community state that lasts for a fixed interval of community biomass *η*; all other species are set to slow growth *r* in that interval. Illustrative example shown in (A) with *K*_*s*_ = 4, *N* = 10 total species and growth dominance *p* = *r*_+_*/r*_*−*_ = 100. After many serial-dilution cycles, some species persist (blue) while others go extinct (red). (B) Average number of surviving species with *N* = 100 exceeds naive expectation *N/K*_*s*_ (black line). (C) The average number of ‘state shares’ for surviving species and extinct species as a function of *K*_*s*_; here, ‘state share’ is the share of community states that each species grows quickly in. For example, Species A in Fig. 5A receives 1/4 of each state in grows quickly as it shares the fast growth with three other species, and thus has a total of 1.25 state shares. (D) *K*_*n*_ model of random preferences: Each species is assigned a high growth rate *r*_+_ in *K*_*n*_ randomly chosen community states and slow growth *r*_*−*_ in other states. Illustrative example shown with *K*_*n*_ = 4, *N* = 10. (E,F) Same as (B,C) but for the *K*_*n*_ model. (F) Average number of ‘state shares’ for surviving species and extinct species as a function of *K*_*s*_; ‘state share’ is defined as in (C), but here each species may share each state with a different number of species. (G) Time course of normalized biomass for the surviving species over a stable cycle in an illustrative example of 10 species with *K*_*s*_ = 4 preferred niches and *K*_*n*_ = 4 competitors in each niche (imposing constraints of both *K*_*s*_ and *K*_*n*_ models). Species show a complex disordered pattern of growth and no-growth states with subtle correlations that enable coexistence. (Error bars in all panels indicate standard deviation from 10 simulations of different growth matrices with 30% variation in *r*_+_, *r*_*−*_, and Δ*η*.)

To gain insight into which species survive, contrast rows of the matrix corresponding to surviving (blue) and extinct species (red) in Fig. 5A. Survivors (blue) tend to grow fast in five or more community states while extinct species (red) have mostly three preferred states. To test this hypothesis, we plotted the average number of preferred states (states in which a species grows at *r*_+_) for the surviving and extinct species for the *N* = 100 system (Fig. 5C); we see that the surviving species are generalists who can grow rapidly in multiple community states.

We next explore the situation where each species has a fixed number *K*_*n*_ of preferred states, i.e., now, all species are generalists to an equal extent. However, the identity of the preferred community states for each species is randomly chosen; see Fig. 5D for an example matrix. The number of surviving species is now clearly increased compared to the *K*_*s*_ model; compare Fig. 5E to Fig. 5B. What factors distinguish the surviving and extinct species? The specific example in Fig. 5D suggests that extinct species had more competitors in their preferred community states. To confirm this, we calculated the average number of preferred states for each species, weighted by the number of competitors that also grow fast in that state. Plotting this competition-weighted average state share for the *N* = 100 system, we find the surviving species (blue) indeed have a higher share of their preferred community states compared to extinct species (red). See Fig. 5F.

In Supplementary Figure S7 and S8 and Supplementary Text S2.9, we consider a Consumer Resource model in a chemostat with each species growing on multiple resources. We explored similar constraints as above, with the total number of species growing on each resource (*K*_*s*_) fixed or the number of resources each species grew on (*K*_*n*_) fixed. We find that the fraction of surviving species for fixed *K*_*n*_ (43% for *K*_*n*_ = 10) is lower than in the CS Model (52% for *K*_*n*_ = 10) and higher for fixed *K*_*s*_ (34% for *K*_*s*_ = 10) than in the CS Model (28% for *K*_*s*_ = 10).

## Discussion

In this work, we have investigated models of growth and survival of microbial species in communities subjected to cyclic environmental fluctuations, focusing on the case of prolonged periods between nutrient replenishment as seen often in the wild [34, 44]; under these conditions, exponential steady-state growth cannot be sustained. In the lab, non-steady-state growth can occur during serial-dilution cycles where the cycle length is long enough for nutrient depletion or build up for toxic waste that limits growth [12, 36, 45]. Accurate bottom-up models are not feasible given our limited understanding of microbial behaviors outside of exponential growth [44, 46].

Inspired by a model experimental system, we investigated the creation of dynamical niches by a combination of physiological states taken by members of a community in response to self-generated environmental changes. In our model, each species *X* is assigned a growth rate *r*_*X*_ (*S*) in each state *S* of the community. A community state *S* was taken to last for an interval of total community biomass *ρ*_*tot*_ accumulated during the growth of the community, and the corresponding growth rate of each species in that community state *S* reflects the physiological state each species is in as well as its environmental context (which includes the physiological states of all other species). Using total community biomass *ρ*_*tot*_ as a driver of community state transitions gives a simple model that allows us to derive quantitative self-consistency conditions on temporal dynamics during serial-dilution cycles using experimentally measurable quantities: Suppose a set of species in repeated serial-dilution cycles are observed to grow at growth rates {*r*_*α*_*}* at time *t* where the community has total biomass *ρ*_tot_(*t*), to what extent can the set of data {*r*_*α*_, *ρ*_tot_} recapitulate the existence of species and the dynamics of their abundances during the cycle? And how robust is the observed dynamics to perturbations in environmental factors and community composition? Most of the results derived in this study are centered around these questions.

One major finding is that community states cannot be taken for granted as “niches” - even when species “take turns” dominating growth in different community states, many species can go extinct. Instead, we find quantitative constraints on how fast or for how wide an interval the dominant species in each community state niche can grow. These constraints can be summarized as: (1) *Tolerance to growth dominance:* increasing the growth rate of an individual species in its favored community state beyond a point does not impact coexistence. This effect contrasts starkly with steady-state coexistence, where the increased growth rate of one species can drive other species extinct. (2) *Community stability through staggered growth dominance:* Stability of a diverse community requires minimizing simultaneous fast growth of multiple species; that is, stability requires that species stagger their growth dominance across distinct community states. (3) *Increased growth dominance for late species:* species growing in late community states must grow faster or for larger biomass intervals than species in early states.

By showing that such niches can arise for a distinct mechanistic reason – transitions between physiological states – the Community State model makes distinct predictions about the relationship between physiology and ecology. Unlike in many other models, niches here are not created by a balance between microbes in exponential growth but originate through an interplay of switches between multiple growing and non-growing states. Our mutual invasibility criteria Eq. **3** offer a quantitative and intuitive understanding of the nature of these niches, factors that widen them, and the nature of competition between species in overlapping niches. The explicit dependence on the ratio of the growth rate of the invading species to that of the resident species in *I*_*A,B*_ and *I*_*B,A*_ make them a fitness-like measure for the current context (cyclic environments with multiple physiological states) where other fitness measures (e.g., growth rate difference) are not applicable.

These effects persist in extensions of the model to many-species communities provided that the number of community states does not greatly exceed the growth dominance *p*, where *p* is the ratio of the growth rate of the dominant species in a community state to the basal growth rate of subordinate species in that state. Trajectories of species abundances (Fig. 5H) show that the dynamics in such systems no longer follow orderly succession dynamics but instead, show a seemingly-disordered array of growth curves that are nevertheless cyclic and hence maintain coexistence. To our knowledge, such disorderly, yet cyclic growth characteristics represent a new class of non-steady-state dynamics that has not been described in the ecology literature.

The tendency in ecology, with an emphasis on steady states, has been to “coarse-grain” or ignore dynamics seen in real systems. Indeed, dynamics in some communities are merely complications (e.g., periodic perturbations about a stable steady state) that can be coarse-grained without any loss of understanding. In fact, in 1973, Stewart and Levin noted mathematically that two species could survive on a single “seasonal resource” [47]. Their work has often been dismissed (including in their own paper [47]) as a mathematical observation relevant only for an assumed growth-affinity trade-off, narrow resource competition, and other idealizations. Our work argues that their simple mechanism of dynamic coexistence – also explored in recent works [41, 48–52] – is *more* relevant, not less, given the observed complex physiology and nonlinear growth dependencies in real microbial ecosystems. If physiological state changes turn out to be dominant drivers of dynamical niches, as seen in [12, 16, 53], dynamics cannot be “averaged” over but become the essential link between physiology and ecology.

We regard an attractive feature of the top-down Community State model to be its direct quantitative connection to experimentally-accessible variables, as well as its avoidance of often inaccessible interaction parameters. Instantaneous growth rates of individual species can be obtained from transient changes in species abundances (via e.g., 16S sequence as proxy), and the total community biomass can be obtained by measuring total protein or total RNA as proxies, or simply by the optical density if the culture does not aggregate. The model studied here, therefore, provides a roadmap for the quantitative analysis of community-wide data to learn about community dynamics, going beyond taxonomic characterization, without invoking fitting parameters. This contrasts starkly with dynamical analysis based on commonly used bottom-up models which invariably involve a large number of unconstrained interaction parameters (e.g., the species interaction matrix in generalized Lotka-Volterra models, or the nutrient consumption matrix in Consumer-Resource models). Additionally, it emphasizes intra-cycle dynamics which has been largely neglected except for a few recent studies [16, 50, 51, 53], and gives concrete predictions, e.g., on the growth rate and duration of early vs late species, that can be tested directly by data. In this sense, the Community State model is a phenomenological model that can be updated directly from data.

Our approach shares common elements with other top-down approaches like the Stochastic Logistic Model [54, 55] and recent data-driven models [56, 57] without explicit interspecies interactions. While these other models attribute growth rate fluctuations to external factors, our model focuses on endogenously-driven environmental change. Our model can be extended to incorporate external fluctuations that randomly perturb growth niches, either across hosts or across cycles, predicting various abundance distributions as in [54]. However, a key distinction is that our approach imposes closure conditions on growth rate variations in repeated cycles needed for stable but dynamic coexistence.

The key idea in our work is the existence of global community states that can be sensed by microbes in that community. Our results suggest that it would be advantageous for organisms to use this information to adjust their behavior and grow in specific community states since such regulation would maximize their chance of survival in the community. For example, organisms occupying early phases of the cycle may benefit from limiting their own growth so as not to eliminate other species active later in the cycle, as late species could be important for the survival of all species in later phases of the cycle – as is the case for acid-induced stress relief [16], the early blooming acid-producing sugar eater is rescued from death by the late-blooming acid consumer which removes the excreted acid and restores the environment.

A more speculative aspect of the Community State model is that the sequence of community states can be parameterized by a one-dimensional eco-coordinate (as opposed to environmental factors or abundances of individual species). A further assumption that allowed for deriving quantitative coexistence criteria and relating them to empirical data is that community biomass can serve as this coordinate parameterizing the sequence of community states. We believe this hypothesis is biologically plausible: First, a number of key physiological parameters, e.g., pH, oxygen content, waste products, and iron availability, change with the accumulation of community biomass [58], and the values of these parameters to cause transitions in the physiological states of individual organisms are known. Other physiological effects such as lag time and cell death might introduce limitations for our framework that require further study [50, 51, 59]. Second, it is known that several autoinducers are produced and sensed by a wide range of both gram-positive and gram-negative bacteria [60–62]. In fact, AI-2 has been proposed to serve as a “universal signal” for inter-species communication [63–65]. Third, it is common for microbes to develop sensors to detect important features of their environment [66–69]; as total community biomass is clearly an important dynamical variable that can be used to forecast the fate of the community (e.g., how close to the carrying capacity), it would not be surprising if organisms have evolved various proxy schemes to sense the total biomass. As bacteria feature multiple sensors and regulatory processes, they may detect various (and possibly distinct from other species) aspects of the global state of the community and integrate the available information through diverse regulatory mechanisms. Thus, community biomass may be viewed as a simplified description to summarize the effects of the different sensors.

The defining feature of the Community State model is the ability of organisms in a community to sense common features of the community and their ability to modulate their own physiology in response to such community-wide signals. Indeed, the existence of a group of organisms that can sense and respond to common features in the environment may be taken as a key characteristic that defines a “community”.

## Acknowledgments

This work was initiated at the Microbial ecology workshop that AM and TH participated in at the NSF-sponsored Kavli Institute of Theoretical Physics (NSF PHY-1748958.). We are grateful for helpful discussions with numerous colleagues during the course of this work, including Milena Chakraverti-Wuerthwein, Otto Cordero, Jacopo Grilli, Akshit Goyal, K.C. Huang, Seppe Kuehn, Pankaj Mehta, Ned Wingreen, and members of the Hwa lab. AVN and TH are supported by the Simons Foundation through the Principles of Microbial Ecosystems (PriME) collaboration (Grant no. 542387) and by the NSF (MCB-2029480). AM is supported by NIGMS of the NIH under award number R35GM151211 and the NSF through the Center for Living Systems (grant no. 2317138).

## Methods

All numerical results in Figs. 1 and 2 were obtained by simulations performed in Matlab (code available at https://github.com/avaneeshnarla/dynamic-metabolic). All other results were obtained using simulations performed in Python 3 using forward Euler integration. For the results in Fig. 4, the integration was performed in time with a growth period of 20 units and a time resolution of 0.001 units. For Fig. 5, the integration was performed in the normalized biomass coordinate (*η*) ranging from 0 to 1, with a resolution of 0.001 units distributed evenly between the niches (such that each niche required the same number of forward integration steps). The initial population abundances in all cases were random fractions of *δ* · 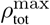, a dilution of the maximum biomass attainable by the population as per our model. Random numbers were drawn from a uniform distribution using Python 3’s Random package.

