## Supplemental Figures and Text for "Dynamic coexistence driven by physiological transitions in microbial communities"

### Supplementary Information for Dynamic coexistence driven by physiological transitions in microbial communities

A Preprint

January 4, 2024

### S1 Supplementary Figures

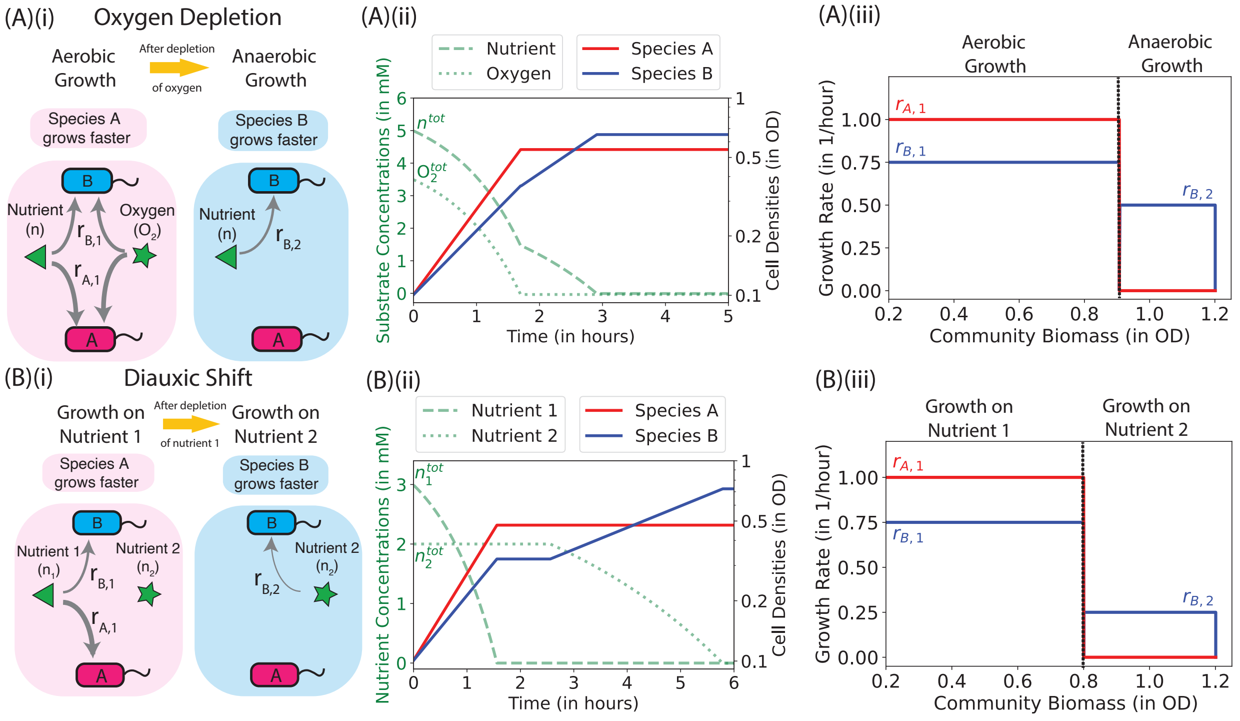

Figure S1: **Growth transitions arising in oxygen depletion and diauxic shift.** Illustrations are made for two species, A (red) and B (blue), whose growth rates, $r_A$ and $r_B$, respectively, vary due to changes in the concentrations of nutrients or toxins in the common media due to a variety of interactions. Growth curves are shown as red and blue solid lines in the middle column, with nutrient/toxin concentrations shown as green dashed or dotted lines. Growth rates are plotted against the accumulated total biomass density in the right column. **A.** Species A grows faster than B aerobically. However, after the depletion of oxygen, only B continues to grow anaerobically, although at a slower rate. **B.** Species A grows faster than B on nutrient 1. Species A stops growing after the depletion of nutrient 1 but B continues to grow on nutrient 2 after a classic diauxic shift, reflected by a lag in the growth curve (middle column of panel B). Note that when plotting the growth rate against the total biomass accumulated in the community, the lag does not show up as there is no biomass accumulation during this period. More such examples are provided in Fig. S2

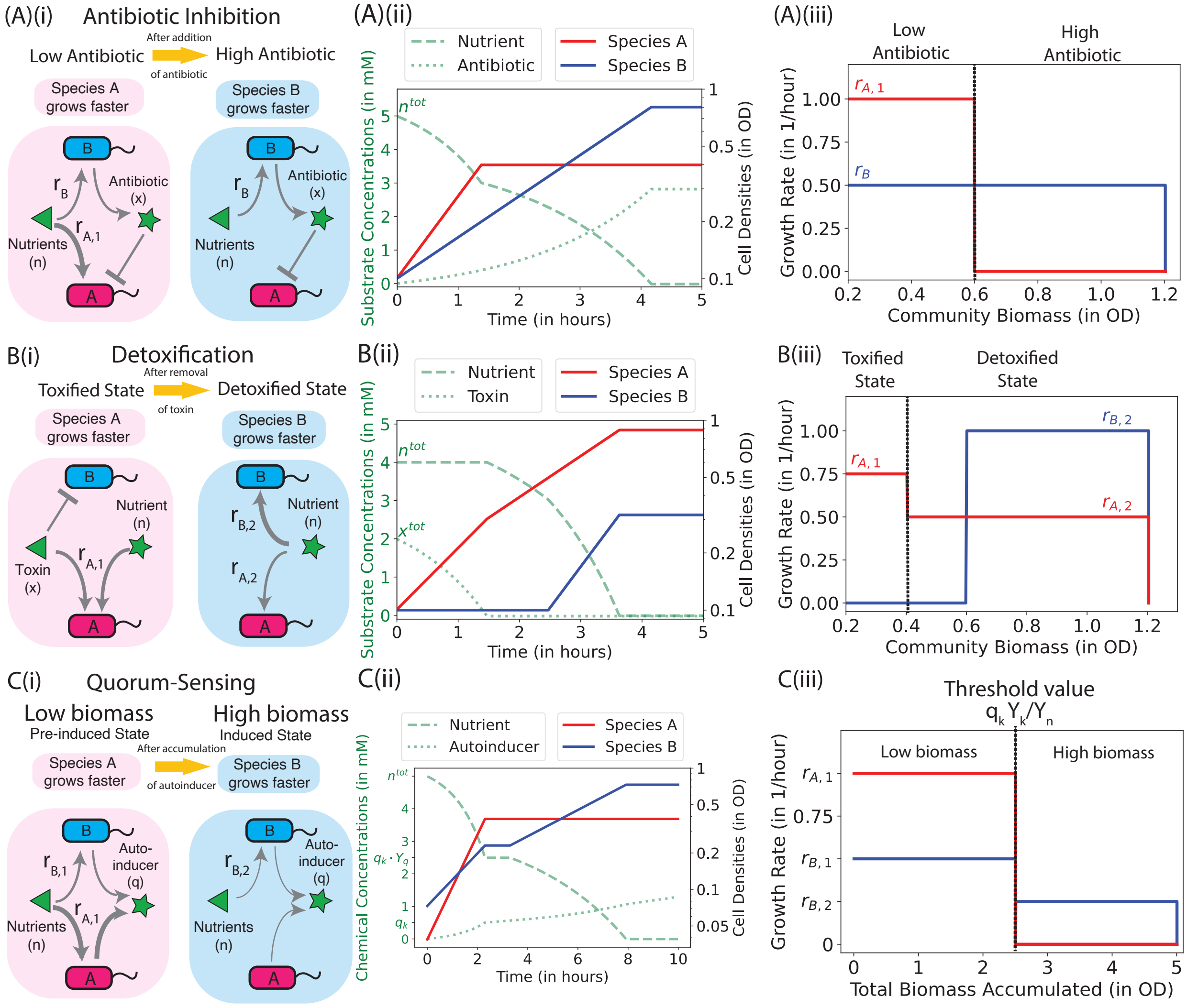

Figure S2: **Growth transitions arising in antibiotic inhibitions, detoxification, and quorum-sensing.** Illustrations are made for two species, A (red) and B (blue), whose growth rates, $r_A$ and $r_B$, respectively, vary due to changes in the concentrations of nutrients or toxins in the common media due to a variety of interactions. Growth curves are shown as red and blue lines in the middle column, with nutrient/toxin concentrations shown as the green dashed or dotted lines. Growth rates are plotted against the accumulated total biomass in the right column. **A.** Species A grows faster than Species B, but Species B excretes an antibiotic which inhibits the growth of Species A when accumulated to a sufficient level. B itself is not sensitive to the antibiotic. **B.** A toxin in the medium inhibits the growth of Species B, but is consumed by Species A. After toxin removal, A continues to grow (at a moderately reduced rate) while B can grow at a fast rate. **C.** Both species secrete autoinducers. After the autoinducer concentration reaches a threshold value, B grows faster than A.

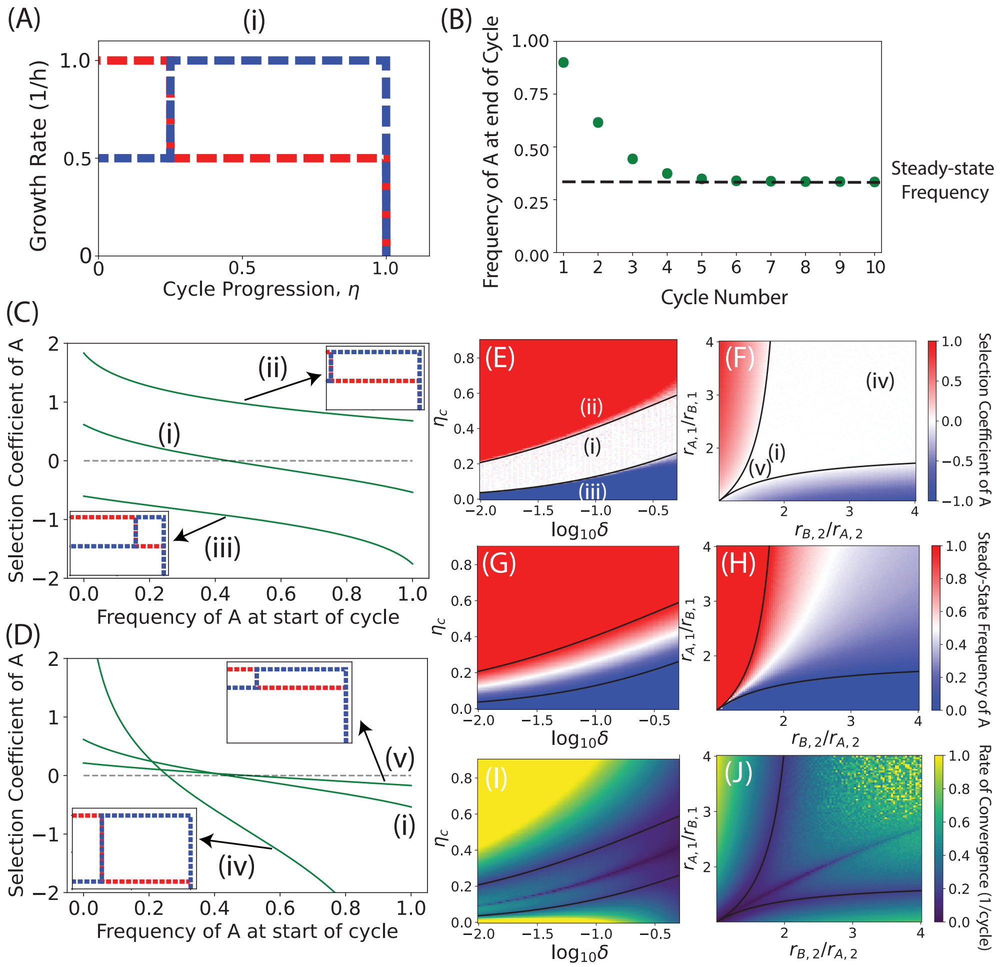

Figure S3: **Partitioning of growth rates reveals quality of coexistence.** Features from growth rate dependences on $\eta$ can be understood by considering perturbations to a piece-wise linear growth rate dependence of two species as shown in (A) and hereby denoted by (i). For all parameters, the coculture eventually arrives at a steady-cycle frequency at the end of the cycle that does not change in subsequent cycles as shown in (B). All results indicated below are for the steady cycle. (C,D) The variation of selection coefficient of A with initial frequency of A for different growth rate dependences (shown in insets and labeled). If the selection coefficient is not 0 for any initial frequency, the two species will not coexist. The effect of the perturbations can be understood more systematically in phase plots where either the environment (in panels E,G,I) or the physiology (in panels F,H,J) of the two species is varied while holding all other parameters constant.

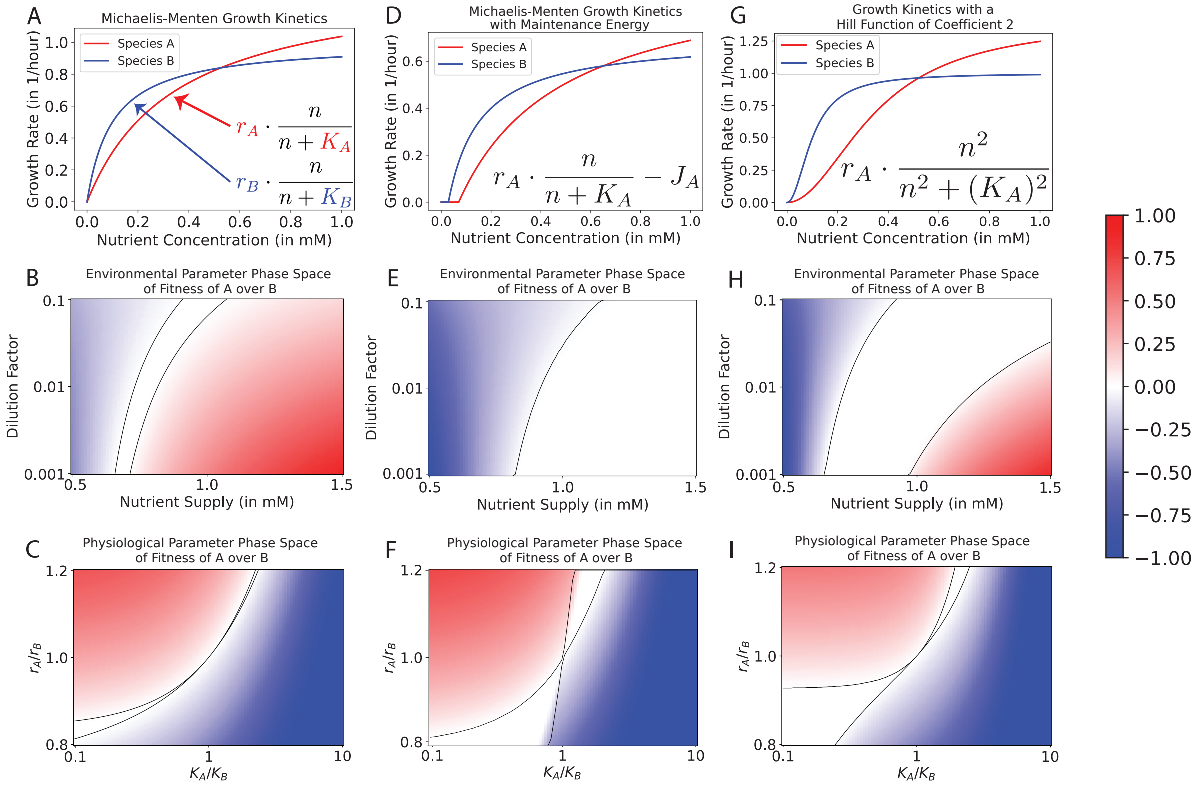

Figure S4: **Biologically-motivated modifications to the Monod growth relation enlarge the coexistence region of phase space.** **A**. Plot showing the standard Monod Growth relation for the dependence of the growth rate on the nutrient concentration of the medium. Two species with different constants describing the growth rates ($r_i$ and $K_i$) are shown and labeled as Species A and B. **B, E, and H**. Plot showing the fitness of species A over species B over one cycle after 100 cycles (common colorbar shown to the right) for different values of $r_A/r_B$ and $K_A/K_B$ for the respective growth functions of each row. The black lines indicate the boundaries of the analytically determined phase boundaries of coexistence. **B**. Plot showing the fitness of species A over species B over one cycle after 100 cycles (common colorbar shown to the right) for different values of environmental parameters (the nutrient supplied at the beginning of each growth cycle and the dilution factor). The black lines indicate the boundaries of the analytically determined phase boundaries of coexistence. **C**. Plot showing the fitness of species A over species B over one cycle after 100 cycles (common colorbar shown to the right) for different values of physiological parameters of species A, shown as ratios $r_A/r_B$ of the two growth rates and $K_A/K_B$ of the two saturation constant. The black lines indicate the boundaries of the analytically determined phase boundaries of coexistence.

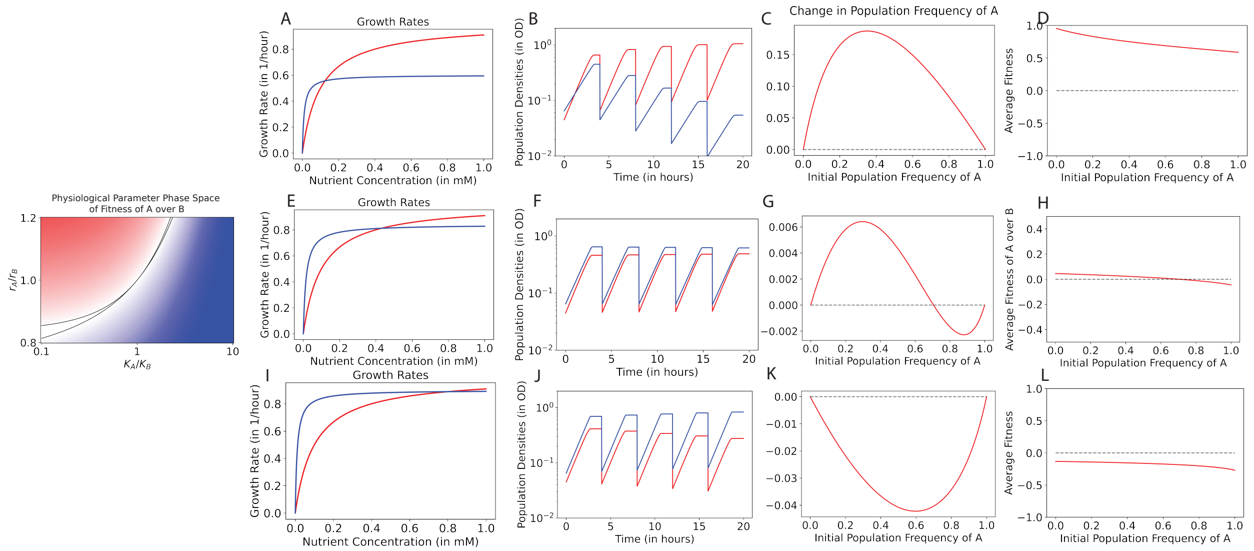

Figure S5: **Assembly of complex communities can be understood by the mutual invasibility criterion for coexistence. A.** The growth functions of two species in competition (either of the species can be replaced by a community with the growth rate indicating the growth rate of the entire community, averaged by the frequency of each member of the community). This is determined entirely by the physiological parameters of the two strains. **B.** The ratios of the two growth rates indicate the differential fitness within the cycle. **C.** The differential fitness needs to be weighted by a resource consumption kernel, given by $\omega(s)$ which is a function of the environmental variables, $s_0$ and $\delta$. If the integral of $r_A(s)/r_B(s)$ curve is $\geq 1$, Species A can invade the monoculture of B, and similarly for A. **D.** Iterative flow maps for both Species A and species showing that if both monocultures are invadible, then one fixed point must exist. The invasibility criteria allows one to infer the existence of a coexistence fixed point. If either monoculture is invadable, the trivial fixed point of that monoculture is unstable. As shown in the text, only one non-trivial fixed point can exist.

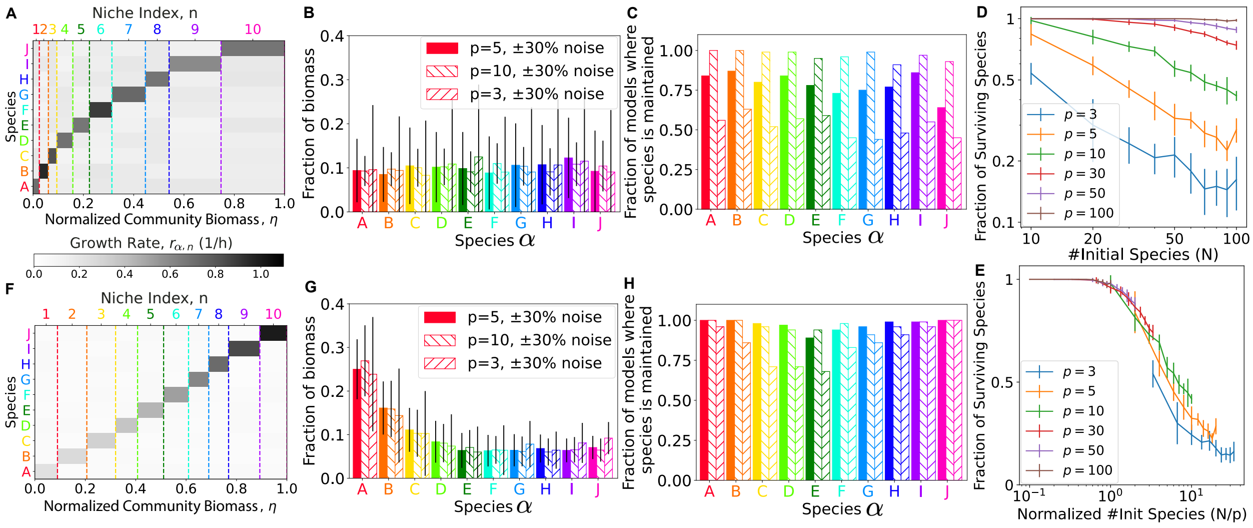

Figure S6: **Impact of disorder and relative changes in growth rate across niches on coexistence** (A) Model with exponential niche width, with the value of growth rate for each diagonal entry (i.e., during the preferred niche) assigned randomly within a range of $r_+$, each off-diagonal entry (non-preferred niches) assigned randomly within a range of $r_-$. (B) The abundance of each species at the end of the stable cycle, obtained for the model parameters indicated, with $p \equiv r_+/r_-$ being the growth preference. The black lines indicate the standard deviation. (C) The fraction of models where each species is maintained, for the same set of parameters as those indicated in the legend of panel B. (D, E) Abundance of the surviving species for different number of niches $N$ and different growth preferences. Panels F, G, H are the same as panels A, B, C, except that niche widths are fixed to a constant, but the growth preference $r_+/r_-$ is varied using Eq. **S123**

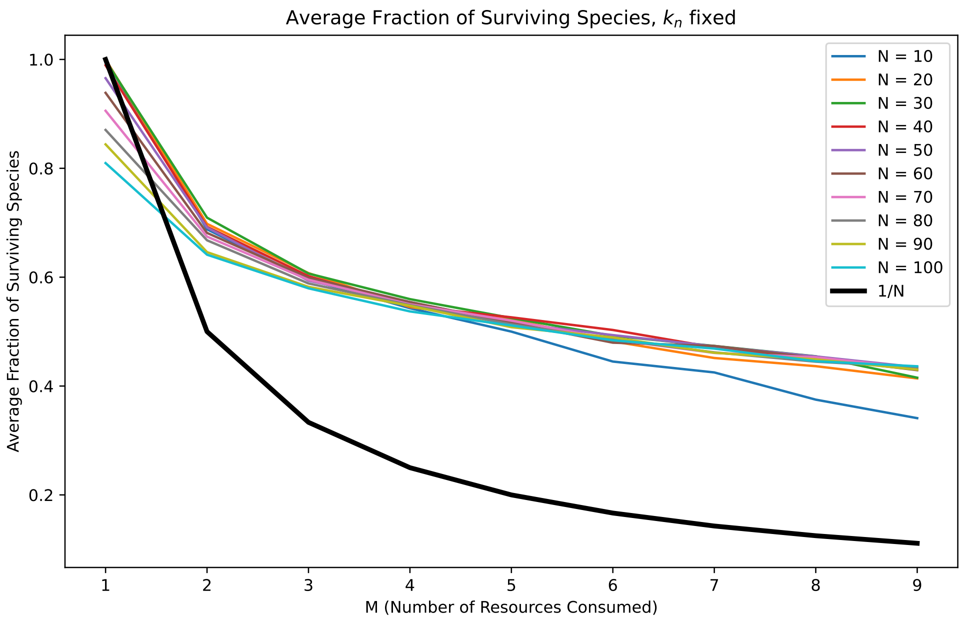

Figure S7: Average Fraction of Surviving Species with $K_n = M$: The plot shows the average fraction of surviving species in a chemostat model where the number of resources consumed per species ($K_n$) is held constant. The survival fractions are plotted against varying numbers of resources consumed per species ($M$). Each line represents a different total number of species/resources ($N$). The solid black line is $1/M$.

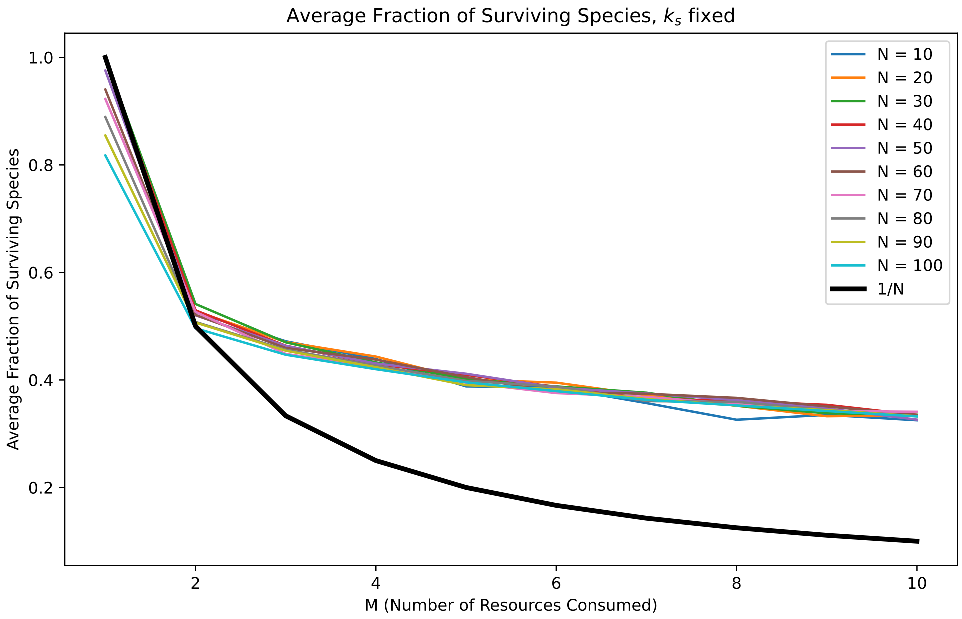

Figure S8: Average Fraction of Surviving Species with $K_s = M$: The plot shows the survival fractions of species under the condition where the number of species consuming each resource ($K_s$) is constant. The plot shows how species survival varies with the change in the number of resources consumed per species ($M$). Different lines correspond to different total numbers of species/resources ($N$). The solid black line is $1/M$.

### S2 Supplementary Text

#### S2.1 Context on the Competitive Exclusion Principle

The Competitive Exclusion Principle (CEP) states that complete competitors cannot coexist [1]. The statement originates with Volterra who used a mathematical model to demonstrate that two species whose growth is limited by the same resource cannot coexist indefinitely [2]. This idea was further explored, developed, and disseminated by Lotka [3], Gause [4, 5], and Hutchinson [6]. Subsequent theoretical work by MacArthur, Levins, and others [7–10] extended the CEP to state that, in general, there can be no more species than resources.

The CEP is also closely tied to the notion of an ecological niche [11, 12] and is alternatively stated as “No two species can indefinitely continue to occupy the same ecological nich” [13]. Levin showed that two species cannot occupy a niche (defined as the hypervolume where the dimensions are environmental conditions and resources, following Hutchinson’s definition [6]) unless their limiting factors (for example, nutrients) differ and are independent [14]. This echoes MacArthur when he says that the proper statement of the CEP is that “species divide up the resources of a community in such a way that each species is limited by a different factor.” [15]. We must note that this holds for both biotic and abiotic factors [2, 7, 9, 14].

All the attempts described above contained the assumption that the specific growth rates of the competing species are linear functions of resource or factor densities [16]. Further, most attempts (with the exception of Levin [14]) only considered coexistence at fixed densities. In fact, it can be shown that coexistence at fixed densities is limited by the number of resources the species can grow on [16], regardless of the form of the growth rates. When both these constraints are simultaneously relaxed, many species can coexist on a few biotic resources [16–18]. A key result was provided by Levins who showed that the number of effective resources is the number of original resources plus the number of distinct non-linearities in the system [19].

Constructive examples rely on the periodic solutions of the Lotka-Volterra model [2] and different points of saturation in the non-linearities of the specific growth rates (we note that nonlinear saturating functional responses are more biologically accurate than linear growth rates [20]). For the case of abiotic resources, Smale showed that that the ordinary differential equation commonly used to describe competing species are compatible with any dynamical behavior provided the number of species is greater than three [21]. Following this, systems have been constructed that have periodic orbits [22] with more than three species. Even chaotic behavior has been observed for the case of essential nutrients [23]. However, such constructed examples require a large number of species and/or nutrients, need to be carefully constructed, and there is no evidence that they occur naturally. Concurrently, in the 1970s, the importance of non-equilibrium interactions of competing populations in establishing species diversity was emphasized in several investigations [24] (See Chapter 15 of [25]).

##### S2.2 Mapping between different mechanistic models for step-wise growth

###### S2.2.1 Base Model

We first describe the base model for the growth step of the growth-dilution cycle, and below we will map different consumer-resource models to the base model mathematically.

We take $\rho_A$ to be the population of Species A, $\rho_B$ to be the population of Species B, and $\eta$ to be the eco-coordinate.

$$\dot{\rho_A} = r_A(\eta) \cdot \rho_A, \tag{S1}$$

where $r_A(\eta) = r_{A,1}$ when $\eta < \eta_k$ and $r_A(s) = r_{A,2}$ when $\eta > \eta_k$.

$$\dot{\rho_B} = r_B(\eta) \cdot \rho_B, \tag{S2}$$

where $r_B(\eta) = r_{B,1}$ when $\eta < \eta_k$ and $r_B(s) = r_{B,2}$ when $\eta > \eta_k$.

$$\dot{\eta} = r_A(\eta) \cdot \rho_A + r_B(\eta) \cdot \rho_B. \tag{S3}$$

$\eta$ starts at 0 and ends at 1.

###### S2.2.2 Step-Wise Growth

We take $N_A$ to be the population of Species A, $N_B$ to be the population of Species B, and $s$ to be the nutrient. The consumer resource model is given by

$$\dot{N_A} = r_A(s)N_A, \tag{S4}$$

where $r_A(s) = r_{A,1}$ when $s > s_k$ and $r_A(s) = r_{A,2}$ when $s < s_k$.

$$\dot{N_B} = r_B(s)N_B, \tag{S5}$$

where $r_B(s) = r_{B,1}$ when $s > s_k$ and $r_B(s) = r_{B,2}$ when $s < s_k$.

$$\dot{s} = -r_A(s)N_A/Y_A - r_B(s)N_B/Y_B. \tag{S6}$$

The environmental variables are given by $s_0$, the total amount of nutrient supplied at first, and $\delta$, the dilution fold.
We redefine the variables as follows:

$$\eta \equiv 1 - \frac{s}{s_0} \tag{S7}$$

$$\eta_k \equiv 1 - \frac{s_k}{s_0} \tag{S8}$$

$$\rho_A \equiv \frac{N_A}{s_0 \cdot Y_A} \tag{S9}$$

$$\rho_B \equiv \frac{N_B}{s_0 \cdot Y_B} \tag{S10}$$

This creates an exact equivalence to the base model.

###### S2.2.3 Diauxie

We take $N_A$ to be the population of Species A, $N_B$ to be the population of Species B, $s_1$ to be the concentration of the first nutrient consumed and $s_2$ to be the concentration of the second nutrient consumed. The consumer resource model is given by

$$\dot{N_A} = r_{A,1}(s_1)N_A + r_{A,2}(s_1, s_2)N_A, \tag{S11}$$

where $r_{A,1}(s_1) = r_{A,1} \cdot \Theta(s_1)$ and $r_{A,1}(s_1, s_2) = r_{A,2} \cdot \Theta(-s_1) \cdot \Theta(s_2)$. $\Theta$ is the Heaviside step-function

$$\dot{N_B} = r_{B,1}(s_1)N_B + r_{B,2}(s_1, s_2)N_B, \tag{S12}$$

where $r_{B,1}(s_1) = r_{B,1} \cdot \Theta(s_1)$ and $r_{B,1}(s_1, s_2) = r_{B,2} \cdot \Theta(-s_1) \cdot \Theta(s_2)$.

$$\dot{s_1} = -r_{A,1}(s_1)\frac{N_A}{Y_A \cdot Y_1} - r_{B,1}(s_1, s_2)\frac{N_B}{Y_B \cdot Y_1}. \tag{S13}$$

$$\dot{s_2} = -r_{A,2}(s_2)\frac{N_A}{Y_A \cdot Y_2} - r_{B,2}(s_1, s_2)\frac{N_B}{Y_B \cdot Y_2}. \tag{S14}$$

The environmental variables are given by $s_0^1$ and $s_0^2$, the total amount of nutrients supplied at first, and $\delta$, the dilution fold.

We redefine the variables as follows:

$$\eta \equiv \begin{cases} 1 - \frac{Y_1 \cdot s_1 + Y_2 \cdot s_0^2}{(Y_1 \cdot s_0^1 + Y_2 \cdot s_0^2)}, s_1 > 0 \\ 1 - \frac{Y_2 \cdot s_2}{(Y_1 \cdot s_0^1 + Y_2 \cdot s_0^2)}, s_1 = 0 \end{cases} \tag{S15}$$

$$\eta_k \equiv 1 - \frac{Y_1 \cdot s_0^1}{(Y_1 \cdot s_0^1 + Y_2 \cdot s_0^2)} \tag{S16}$$

$$\rho_A \equiv \frac{N_A}{Y_A \cdot (Y_1 \cdot s_0^1 + Y_2 \cdot s_0^2)} \tag{S17}$$

$$\rho_B \equiv \frac{N_B}{Y_B \cdot (Y_1 \cdot s_0^1 + Y_2 \cdot s_0^2)} \tag{S18}$$

This gives us that:

$$\dot{\rho_A} = r_A(\eta)\rho_A, \tag{S19}$$

where $r_A(\eta) = r_{A,1} \cdot \Theta(\eta_k - \eta)$ and $r_A(\eta) = r_{A,2} \cdot \Theta(\eta - \eta_k) \cdot \Theta(1 - \eta)$.

$$\dot{\rho_B} = r_B(\eta)\rho_B, \tag{S20}$$

where $r_B(\eta) = r_{B,1} \cdot \Theta(\eta_k - \eta)$ and $r_B(\eta) = r_{B,2} \cdot \Theta(\eta - \eta_k) \cdot \Theta(1 - \eta)$.

$$\dot{\eta} = r_A(\eta)\rho_A + r_B(\eta)\rho_B. \tag{S21}$$

This creates an exact equivalence to the base model.

###### S2.2.4 Oxygen Depletion

We take $N_A$ to be the population of Species A, $N_B$ to be the population of Species B, $s$ to be the concentration of the nutrient, and $x$ to be the concentration of oxygen. The consumer resource model is given by

$$\dot{N_A} = r_A(s, x)N_A, \tag{S22}$$

where $r_A(s, x) = r_{A,1} \cdot \Theta(x) + r_{A,2} \cdot \Theta(-x) \cdot \Theta(s)$.

$$\dot{N_B} = r_B(s, x)N_B, \tag{S23}$$

where $r_B(s, x) = r_{B,1} \cdot \Theta(x) + r_{B,2} \cdot \Theta(-x) \cdot \Theta(s)$.

$$\dot{s} = -r_A(s, x)N_A/Y_A - r_B(s, x)N_B/Y_B. \tag{S24}$$

$$\dot{x} = (-r_{A,1} \cdot \frac{N_A}{Y_A \cdot Y_x} - r_{B,1} \cdot \frac{N_B}{Y_B \cdot Y_x})\Theta(x). \tag{S25}$$

The environmental variables are given by $s_0$ and $x_0$, the total amount of nutrient and oxygen supplied at first, and $\delta$, the dilution fold. For oxygen to deplete first, we need $x_0 < s_0 \cdot Y_x$

We redefine the variables as follows:

$$\eta \equiv 1 - \frac{s}{s_0} \tag{S26}$$

$$\eta_k \equiv 1 - \frac{x_0}{s_0 \cdot Y_x} \tag{S27}$$

$$\rho_A \equiv \frac{N_A}{s_0 \cdot Y_A} \tag{S28}$$

$$\rho_B \equiv \frac{N_B}{s_0 \cdot Y_B} \tag{S29}$$

This creates an exact equivalence to the base model.

###### S2.2.5 Quorum Sensing

We take $N_A$ to be the population of Species A, $N_B$ to be the population of Species B, $s$ to be the concentration of the nutrient, and $q$ to be the concentration of autoinducer. The consumer resource model is given by

$$\dot{N_A} = r_A(s,x)N_A, \tag{S30}$$

where $r_A(s,x) = r_{A,1}$ when $q < q_k$, $r_{A,2}$ when $q > q_k$, and 0 when $s = 0$.

$$\dot{N_B} = r_B(s,x)N_B, \tag{S31}$$

where $r_B(s,x) = r_{B,1}$ when $q < q_k$, $r_{B,2}$ when $q > q_k$, and 0 when $s = 0$.

$$\dot{s} = -r_A(s,x)N_A/Y_A - r_B(s,x)N_B/Y_B. \tag{S32}$$

$$\dot{q} = r_{A,1} \cdot \frac{N_A}{Y_A \cdot Y_q} + r_{B,1} \cdot \frac{N_B}{Y_B \cdot Y_q}. \tag{S33}$$

The environmental variables are given by $s_0$, the total amount of nutrient supplied at first, and $\delta$, the dilution fold. For the autoinducer to accumulate first, we need $q_k < s_0 \cdot Y_q$
We redefine the variables as follows:

$$\eta \equiv 1 - \frac{s}{s_0} \tag{S34}$$

$$\eta_k \equiv 1 - \frac{q_k}{s_0 \cdot Y_q} \tag{S35}$$

$$\rho_A \equiv \frac{N_A}{s_0 \cdot Y_A} \tag{S36}$$

$$\rho_B \equiv \frac{N_B}{s_0 \cdot Y_B} \tag{S37}$$

This creates an exact equivalence to the base model.

###### S2.2.6 Acid Stress

We take $N_A$ to be the population of Species A, $N_B$ to be the population of Species B, $s$ to be the concentration of the nutrient, $x$ to be the concentration of acid, and $p$ to be the concentration of pyruvate. The consumer resource model is given by

$$\dot{N_A} = r_A(s,x,p) \cdot N_A, \tag{S38}$$

where $r_A(s,x) = r_{A,1}$ when $x < x_c$, $r_{A,2}$ when $x > x_c$, and 0 when $s = 0$.

$$\dot{N_B} = r_B(s,x,p) \cdot N_B, \tag{S39}$$

where $r_B(s,x) = r_{B,1}$ when $x < x_c$, $r_{B,2}$ when $x > x_c$, and 0 when $s = 0$.

$$\dot{s} = -r_A \cdot \frac{N_A}{Y_A \cdot Y_s}. \tag{S40}$$

$$\dot{x} = r_A \cdot \frac{N_A}{Y_A \cdot Y_x} \cdot f_x - r_B \cdot \frac{N_B}{Y_B \cdot Y_x}. \tag{S41}$$

$$\dot{p} = r_A \cdot \frac{N_A}{Y_A \cdot Y_p} \cdot f_p - r_B \cdot \frac{N_B}{Y_B \cdot Y_p}. \tag{S42}$$

such that $f_x + f_p < 1$ and the fractions of the nutrient consumed that Species A converts to acid and pyruvate. The environmental variables are given by $s_0$, the total amount of nutrient supplied at first, and $\delta$, the dilution fold. We redefine the variables as follows:

$$\eta \equiv 1 - \frac{Y_s \cdot s + Y_x \cdot x + Y_p \cdot p}{Y_s \cdot s_0} \tag{S43}$$

$$\rho_A \equiv \frac{N_A}{s_0 \cdot Y_A} \tag{S44}$$

$$\rho_B \equiv \frac{N_B}{s_0 \cdot Y_B} \tag{S45}$$

All that remains to resolve is the question of $\eta_k$ since in the actual system, it is only given by the acetate concentration, $x_c$. However, we can take $x_c$ to be a function of $\eta$ if $s(t) \propto x(t)$:

$$\eta(x = x_c) \equiv 1 - \frac{Y_s \cdot s(x_c) + Y_x \cdot x_c}{Y_s \cdot s_0} \tag{S46}$$

In reality,

$$s(t) = N_A \cdot (\exp(r_{A,1} \cdot t) - 1) \cdot \frac{1}{Y_A \cdot Y_s} \tag{S47}$$

$$x(t) = N_A \cdot (\exp(r_{A,1} \cdot t) - 1) \cdot \frac{1}{Y_A \cdot Y_x} - N_B \cdot (\exp(r_{B,1} \cdot t) - 1) \cdot \frac{1}{Y_B \cdot Y_x} \tag{S48}$$

$$= s(t) \cdot \frac{Y_s}{Y_x} - \frac{N_B}{Y_B \cdot Y_x} \cdot (\exp(r_{B,1} \cdot t) - 1) \tag{S49}$$

$$= s(t) \cdot \frac{Y_s}{Y_x} - \frac{N_B}{Y_B \cdot Y_x} \cdot \left(\frac{Y_A \cdot Y_s \cdot s(t)}{N_A}\right)^{r_{B,1}/r_{A,1}} \tag{S50}$$

Thus, $s(t) \propto x(t)$ if $r_{B,1} \ll r_{A,1}$ (or $N_B \ll N_A$). This creates an exact equivalence to the base model.

##### S2.3 Rescaling

Here, we discuss the rescaling of the time spent in each phase/niche and the transition points in the piece-wise linear models discussed in the paper. The rescaling of the transition points amounts to a redefinition of the units with $\rho_{\text{tot}}^{\text{max}}$ declared to be 1, with an appropriate normalization of all transition points by $\rho_{\text{tot}}^{\text{max}}$. For the many species model, we additionally define a parameter, $\eta \equiv (\rho_{\text{tot}}/\rho_{\text{tot}}^{\text{max}} - \delta)/(1-\delta)$, for simplification of the results such that the cycle starts at $\eta = 0$ and ends at $\eta = 1$. For rescaling the time spent in each phase, we first note that in the limit of large time between dilution events, there is no activity once the system reaches the total biomass density of the system reaches $\rho_{\text{tot}}^{\text{max}}$. Thus, we may ignore the timescales associated with dilution. Next, we consider the dynamics within niche/phase, $n$:

$$\frac{d}{dt}\rho_\alpha^{(j)} = r_{\alpha,n} \cdot \rho_\alpha^{(j)}(t) \qquad \text{for } \eta_{n-1} \leq \eta \leq \eta_{n-1}. \tag{S51}$$

is essentially independent of every other phase as the dynamical system described by Eq. **1** is autonomous in time. Thus we may define, for example, the dimensionless time, $\tau_n \equiv t \cdot \langle \alpha r_{\alpha,n} \rangle_{\alpha \neq \alpha(n)}$. Hence, rescaling $t$ by $\tau_n$, we obtain that the dynamics of $\rho_\alpha^{(j)}$ in niche $n$ given by

$$\frac{d}{d\tau_n}\rho_\alpha^{(j)} = \frac{r_{\alpha,n}}{\langle \alpha r_{\alpha,n} \rangle_{\alpha \neq \alpha(n)}} \cdot \rho_\alpha^{(j)}(t) \qquad \text{for } \eta_{n-1} \leq \eta \leq \eta_{n-1}. \tag{S52}$$

This allows us to define $p_n \equiv \frac{r_{\alpha,n}}{\langle \alpha r_{\alpha,n} \rangle_{\alpha \neq \alpha(n)}}$. Thus, in the cases without noise described in this manuscript, each niche is effectively described by one number: the domination of the fast-growing species in niche $n$ over the slow-growing species.

##### S2.4 Mathematical Results for the Simple Toy Model

Though we have shown that two species may coexist in growth-dilution cycles, understanding the dynamics of the co-culture system mathematically can be significantly difficult because of the non-linear growth rates involved. To simplify the system, we consider a piece-wise linear approximation of the system (as shown in Fig. 3A). In this Toy Model, we take Species A to grow at a constant rate $r_{A,1}$ and Species B at $r_{B,1}$ while biomass values are above a threshold $s_k$ (we call this the first phase and call the time taken to complete it $\tau_1$). And in the 2nd phase (biomass from 0 to $\eta_k$, taking time $\tau_2$), we take Species A to grow at $r_{A,2}$ and Species B at $r_{B,2}$ (this toy model is described in Fig. 3A). When biomass reaches $\rho_{\text{tot}}^{\text{max}}$, we assume that both species stop growing.

If two species coexist in such a system, it implies that the net growth rate is the same over the steady-state cycle, i.e.,

$$r_{A,1}\tau_1 + r_{A,2}\tau_2 = r_{B,1}\tau_1 + r_{B,2}\tau_2 = -\log\delta. \tag{S53}$$

We call the time taken to complete the first phase, $\tau_1$, and the time taken to complete the second phase, $\tau_2$. If two species coexist in such a system, it implies that the net growth rate is the same over the steady-state cycle, i.e.,

$$r_{A,1}\tau_1 + r_{A,2}\tau_2 = r_{B,1}\tau_1 + r_{B,2}\tau_2 = -\log\delta. \tag{S54}$$

For our purposes, we assume that the time between dilution steps, $T$, is longer than $\tau_1 + \tau_2$. We note that as long as there is a trade-off in the growth rates (such that neither species is growing faster than the other species at all biomass values), there exists a positive $\tau_1$ and $\tau_2$ that can solve Eq. **S53**. However, Eq. **S53** is also coupled with a set of equations describing the biomass increase that feature the populations of both species, $\rho_A(0)$ and $\rho_B(0)$, at the beginning of the cycle:

$$\rho_{\text{tot,c}} - \delta \cdot \rho_{\text{tot}}^{\text{max}} = (e^{r_{A,1}\tau_1} - 1)\rho_A(0) + (e^{r_{B,1}\tau_1} - 1)\rho_B(0), \tag{S55}$$

$$\rho_{\text{tot}}^{\text{max}} - \rho_{\text{tot,c}} = (e^{r_{A,2}\tau_2} - 1)e^{r_{A,1}\tau_1}\rho_A(0) + (e^{r_{B,2}\tau_2} - 1)e^{r_{B,1}\tau_1}\rho_B(0). \tag{S56}$$

We obtain the equations above as the total amount of biomass produced by both species must be $\rho_{\text{tot,c}} - \delta \cdot \rho_{\text{tot}}^{\text{max}}$ in the first phase (as the co-culture starts with biomass of $\delta \cdot \rho_{\text{tot}}^{\text{max}}$ after dilution by a factor of $\delta$ from the maximal biomass value of $\rho_{\text{tot}}$, and $\rho_{\text{tot}}^{\text{max}} - \rho_{\text{tot,c}}$ in the second phase. The solution to Eq. **S55**-Eq. **S56**, however, may not yield a viable positive solution for $\rho_A(0)$ and $\rho_B(0)$. For viable solutions, we require that the two species must necessarily engage in resource sharing in the steady-state cycle described by $\tau_1$ and $\tau_2$ that solve Eq. **S53**, as otherwise they would not be able to accumulate all of the biomass by themselves. If any resources remain unconsumed in a monoculture (say of Species A), a very small population of Species B can consume the remaining resources and grow more than the dilution fold. Eventually, the very small population grows larger and the factor of growth decreases until both species grow at the dilution fold.

Thus, to get $\rho_A(0) > 0$, we must have (assuming $r_{A,1} > r_{B,1} > 0,\ r_{B,2} > r_{A,2} > 0$):

$$\rho_A(0) > 0 \iff \underbrace{\rho_{\text{tot,c}} - \delta \cdot \rho_{\text{tot}}^{\text{max}}}_{\text{Biomass produced in phase 1}} > \underbrace{(e^{r_{B,1}\tau_1} - 1)}_{\text{Net Growth of B in } \tau_1} \cdot \underbrace{\delta\rho_{\text{tot}}^{\text{max}}}_{\text{Max population of B at start of phase 1}} \tag{S57}$$

And similarly, requiring that $\rho_B > 0$, we have that

$$\rho_B(0) > 0 \iff \underbrace{\rho_{\text{tot,c}} - \delta \cdot \rho_{\text{tot}}^{\text{max}}}_{\text{Biomass produced in phase 1}} < \underbrace{(e^{r_{A,1}\tau_1} - 1)}_{\text{Net Growth of A in } \tau_1} \cdot \underbrace{\delta\rho_{\text{tot}}^{\text{max}}}_{\text{Max population of A at start of phase 1}} \tag{S58}$$

Or by looking at phase 2,

$$\rho_B(0) > 0 \iff \underbrace{\rho_{\text{tot}}^{\text{max}} - \rho_{\text{tot,c}}}_{\text{Biomass produced in phase 2}} > \underbrace{(e^{r_{A,2}\tau_2} - 1)}_{\text{Net Growth of A in } \tau_2} \cdot \underbrace{\delta\rho_{\text{tot}}^{\text{max}} \cdot e^{r_{A,1}\tau_1}}_{\text{Max population of A at start of phase 2}} \tag{S59}$$

But Eq. **S57** and Eq. **S58** are just saying that resource sharing must be possible for the necessary $\tau_1$ as we require that the resources consumed by the co-culture in $\tau_1$ be greater than the resources that a monoculture of $B$ would have consumed, and less what a monoculture of $A$ would have consumed (we need the second requirement as otherwise you do reach phase 2 before $\tau_1$, and thus you cannot have coexistence). Similarly, we get that the resources in phase 2 should be more than the monoculture of A could consume on its own. In fact, if we add up the two conditions, we get that you need resource sharing of total biomass must be possible). We note that both Eq. **S57** and Eq. **S58** are not necessarily true (trivially, we can move $\rho_{\text{tot}}^{\text{max}} - \rho_{\text{tot,c}}$ so that either Eq. **S57** or Eq. **S58** are not satisfied as the RHS is independent of $\rho_{\text{tot}}^{\text{max}} - \rho_{\text{tot,c}}$). Thus, only certain monocultures permit resource sharing. Further, we note that the ones that do, are stable since if there's resources that could be consumed that a monoculture cannot consume, it allows for invasion that can grow in a time period given by matrix inversion.

We now solve for $\rho_A(0)$ and $\rho_B(0)$ that satisfy Eq. **S53-S56**. First, from Eq. **S53**, we obtain,

$$\tau_1 = -\log\delta \cdot \left(\frac{r_{A,2} - r_{B,2}}{r_{B,1} \cdot r_{A,2} - r_{B,2} \cdot r_{A,1}}\right) \tag{S60}$$

$$= \log(1/\delta) \cdot \frac{1}{r_{B,1}} \left(\frac{p_B - 1}{p_A \cdot p_B - 1}\right). \tag{S61}$$

And similarly, we obtain

$$\tau_2 = -\log\delta \cdot \left(\frac{r_{A,1} - r_{B,1}}{r_{A,1} \cdot r_{B,2} - r_{A,2} \cdot r_{B,1}}\right) = \log(1/\delta) \cdot \frac{1}{r_{A,2}} \left(\frac{p_A - 1}{p_A \cdot p_B - 1}\right). \tag{S62}$$

We note that this value is independent of $\rho_{\text{tot,c}}$ and $\rho_{\text{tot}}^{\text{max}}$. To obtain the steady-state values of $\rho_A(0)$ and $\rho_B(0)$, which we denote with $\rho_A^*$ and $\rho_B^*$ (note that $\rho_A^* + \rho_B^* = \delta \cdot \rho_{\text{tot}}^{\text{max}}$), we use Eq. **S55-S56** to obatin

$$\rho_{\text{tot,c}} - \delta \cdot \rho_{\text{tot}}^{\text{max}} = (e^{r_{A,1}\tau_1} - 1)\rho_A^* + (e^{r_{B,1}\tau_1} - 1)(\delta\rho_{\text{tot}}^{\text{max}} - \rho_A^*) \tag{S63}$$

$$\implies \rho_A^* = \frac{\rho_{\text{tot,c}} - e^{r_{B,1}\tau_1}\delta\rho_{\text{tot}}^{\text{max}}}{e^{r_{A,1}\tau_1} - e^{r_{B,1}\tau_1}} \tag{S64}$$

$$= \frac{\eta_c - \delta^{\frac{p_A - 1}{p_A - 1/p_B}}}{\delta^{\frac{1-p_B}{p_B - 1/p_A}} - \delta^{\frac{1-p_B}{p_A \cdot p_B - 1}}} \cdot \rho_{\text{tot}}^{\text{max}}, \tag{S65}$$

and similarly,

$$\rho_B^* = \frac{\rho_{\text{tot,c}} - e^{r_{A,1}\tau_1}\delta\rho_{\text{tot}}^{\text{max}}}{e^{r_{B,1}\tau_1} - e^{r_{A,1}\tau_1}} \tag{S66}$$

$$= \frac{\delta^{\frac{p_A - 1}{p_A \cdot p_B - 1}} - \eta_c}{\delta^{\frac{1-p_B}{p_B - 1/p_A}} - \delta^{\frac{1-p_B}{p_A \cdot p_B - 1}}} \cdot \rho_{\text{tot}}^{\text{max}}. \tag{S67}$$

Thus, while there is a non-trivial dependence on $\delta$, $p_A$, and $p_B$, there is a linear dependence on $\eta_c$ and $\rho_{\text{tot}}^{\text{max}}$. Below, we explore the non-trivial dependence on $p_A$, and $p_B$. This analysis also tells us that there can only be one viable non-trivial solution for $\rho_A^*$ and $\rho_B^*$ (in other words, there can only be one non-trivial fixed point for the discrete map given by one growth-dilution cycle). This is why the complete system at steady-state can be understood by studying the trivial fixed points (i.e., the respective monocultures) as if both trivial fixed points are unstable to invasion, the system must have a non-trivial fixed point. Further, we can show that if it exists, the non-trivial fixed point must be stable. Intuitively, this is because if the frequency of Species A is increased, the amount of time it would take for the population to produce all of the allocated biomass in its preferred phase will be shorter, thus harming Species A. Similarly, the amount of time it would take for the population to accumulate all of the biomass in the phase that it is growing slower will be longer, once again harming Species A. Thus, there can only be one stable fixed point in the system.

We note that this method can be extended to many species and many phases as well. While it can be difficult to infer the final solution as it requires solving a system of transcendental equations of many variables, this approach does tell us that there can be exactly one solution for $\{\rho_\alpha^*\}$. This is because there is an unknown variable for each species ($\rho_\alpha^*$), and a corresponding equation (similar to Eq. **S53**) for each species. While this system of equations for each species features an unknown variable for each phase ($\tau_i$), the equations for resource constraints provide another set of equations for each $\tau_i$ (which is the transcendental system of equations). Thus, unless the system of transcendental equations produces a degeneracy for $\tau_i$, there is exactly one $\{\rho_\alpha^*\}$ that solves the system of equations.

##### S2.5 Negative frequency-dependent selection in growth-dilution cycles

Here, we discuss the stabilizing mechanism for coexistence in growth-dilution cycles. We may understand this by considering only the dynamics of frequency of one species, say Species A, denoted by $x(t)$:

$$x(t) \equiv \frac{\rho_A(t)}{\rho_A(t) + \rho_B(t)}. \tag{S68}$$

By changing variables in equations, Eq. **1**, we arrive at the following equation for the dynamics of $x$:

$$\dot{x} = (r_A - r_B) \cdot x \cdot (1 - x). \tag{S69}$$

This is the logistic growth equation with time-varying growth rates. This gives us that the frequency of Species A at the end of the cycle, $x(T)$, given that its frequency at the beginning of the cycle is $x_0$, is

$$x(T) = \frac{x_0}{x_0 + (1 - x_0)e^{-q(x_0)}}, \tag{S70}$$

where $q(x_0)$ is the average fitness of Species A over Species B over one cycle:

$$q(x_0) = \int_0^T (r_A(s(t)) - r_B(s(t)))dt. \tag{S71}$$

This shows that $x(T)$ is uniquely determined by $x_0$ as $s(t)$ can be uniquely determined by solving the initial value problem given in Eq. **1**. Further, from Eq. **S70** we note that the change in frequency of A, $x(T) - x_0$, has the same sign as $q(x_0)$, and $x(T) = x_0 \equiv x^*$ if $q(x^*) = 0$. This just reiterates that the frequency of A will be determined by its average relative fitness over one cycle. Such a point, $x^*$ would be a fixed point of the map that describes how the frequency changes over one cycle, $x_0 \to x(T)$, since for $x = x^*$, the frequency doesn't change.

Thus, the dynamical behavior of the growth-dilution cycle is determined by $q(x_0)$. While determining $q$ is very difficult as we cannot know $s(t)$ without solving the dynamical system given by Eq. **1**, we can show that $q' < 0$ for all $x_0$ (Section S2.5.2). This shows that $q = 0$ for at most one value of $x_0$ and allows us to construct a Lyapunov function,

$$V(x) = \int_x^{x^*} q(x')dx'. \tag{S72}$$

If no $x^*$ exists such that $q(x^*) = 0$, $x^*$ can be taken to the value closest to 0 (which is necessarily unique as $q' < 0$). As verified in Section S2.5.2, $V(x)$ fits all the requisite criteria for a discrete-time strict Lyapunov function that leads to global asymptotic stability. Thus, for any two growth dependences, the system always relaxes to the same steady state.

This result that $q' < 0$ can be restated as saying that the average fitness of a species has a negative frequency dependence in growth-dilution cycles. This result holds for all $r_A$ and $r_B$. Further, the magnitude of $q'$ reveals how stable the system is. If $q'$ is very large, then the system quickly converges to the fixed point (Fig. 3G). A greater negative value of $q'$ indicates that the difference in growth rates is much larger than the mean growth rate. We note that this is the case when the growth dependences are highly non-linear.

##### S2.5.1 Stabilization of Resource Trajectories

A different perspective on the stabilizing nature of resource sharing can be obtained by looking at the trajectory that the co-culture traverses in the space of environmental variables. The key feature in the consumer-resource models that we use is that the trajectory is determined by the growth of the species. This is because the populations affect the environment through growth (either by consuming resources or secreting pollutants). Let us consider perturbations to the resource trajectory corresponding to a fixed point. Higher growth during a section of the trajectory means faster movement along the trajectory and less time spent at that section. But growth is also proportional to the time spent along the resource trajectory. Conversely, slower growth means more time at that section and thus the initial perturbation is countered. This leads to negative feedback for growth and, as a result, for resource change, thus stabilizing the resource trajectory. For the case of many environmental variables, movement along the trajectory can be projected onto the axis corresponding to each environmental variable, and the trajectory along each environmental variable can be considered independently. As movement along each projection is stabilized independently (such that at every point in time, there is a unique stable value of each environmental coordinate), the entire trajectory is stabilized.

A similar understanding can be obtained by considering perturbations in the resource trajectory itself rather than perturbations in growth as we did above. A perturbation in the resource trajectory will either slow down or speed up growth, and this perturbation in growth will counteract the original perturbation in the resource trajectory.

However, this stabilization is local along every point on the resource trajectory, while coexistence is a global property of the entire trajectory. In other words, a negative frequency dependence of fitness does not mean that fitness ever has to be negative. Coexistence requires that each species have a negative fitness over the other species for some frequency, especially when it is abundant. This leads us to a simple necessary and sufficient criterion for coexistence: mutual invasibility (discussed in Section S2.6.

##### S2.5.2 Proof of Negative Frequency Dependence

Consider the frequency of Species A,

$$x_A = \frac{\rho_A}{\sum_\alpha \rho_\alpha} \tag{S73}$$

Taking the time derivative, we have that

$$\dot{x_A} = \frac{\dot{\rho_A} \cdot \sum_\alpha \rho_\alpha - \rho_A \cdot \sum_\alpha \dot{\rho_\alpha}}{(\sum_\alpha \rho_\alpha)^2} \tag{S74}$$

$$= \frac{r_A \cdot \rho_A \cdot \sum_\alpha \rho_\alpha - \rho_A \cdot \sum_\alpha r_A \rho_\alpha}{(\sum_\alpha \rho_\alpha)^2} \tag{S75}$$

$$= \frac{r_A \cdot \rho_A \cdot \sum_{\alpha \neq A} \rho_\alpha - \rho_A \cdot \sum_{\alpha \neq A} r_\alpha \rho_\alpha}{(\sum_\alpha \rho_\alpha)^2} \tag{S76}$$

$$= \frac{r_A \cdot \rho_A \cdot \sum_{\alpha \neq A} \rho_\alpha - \bar{r} \cdot \rho_A \cdot \sum_{\alpha \neq A} \rho_\alpha}{(\sum_\alpha \rho_\alpha)^2} \tag{S77}$$

$$\text{where } \bar{r} \equiv \frac{\sum_{\alpha \neq A} r_\alpha \rho_\alpha}{\sum_{\alpha \neq A} \rho_\alpha} = \frac{\sum_{\alpha \neq A} r_\alpha \cdot x_\alpha}{1 - x_A}. \tag{S78}$$

$$= (r_A - \bar{r}) \cdot x_A \cdot (1 - x_A) \tag{S79}$$

Thus, for any time $t$, we have that

$$x_A(t) = \frac{x_A^0}{x_A^0 + (1 - x_A^0) e^{-q(x_A^0, t)}} \tag{S80}$$

where $x_A^0 = x(t=0)$ and

$$q_A(\{x_A^0\}, t) = \int_0^t (r_A - \bar{r})dt \tag{S81}$$

This gives us that

$$x_A(T) = \frac{x_A^0}{x_A^0 + (1 - x_A^0)e^{-q_A(x_A^0, T)}}. \tag{S82}$$

Since this calculation holds for all species, we have a map from the species population at the beginning of the cycle to the species population at the end of the cycle. We define the Lyapunov function, $V$, such that

$$V_A(\{x_\alpha(0)\}) \equiv \int_{\{x_\alpha(0)\}}^{\{x_\alpha^*\}} q_A(\{x_A\}, T) \cdot d(\{x_A\}). \tag{S83}$$

For discrete time dynamics, the requirement for global convergence to $\{x_\alpha^*\}$ is that $V_A(\{x_\alpha^*\}) = 0$ (by definition), $V_A(\{x_\alpha(0)\}) > 0$, and $V_A(\{x_\alpha(T)\}) - V_A(\{x_\alpha(0)\}) < 0$ [26]. We note that

$$V_A(\{x_\alpha(T)\}) - V_A(\{x_\alpha(0)\}) = -\int_{\{x_\alpha(0)\}}^{\{x_\alpha(T)\}} q_A(\{x_A\}, T) \cdot d(\{x_A\}) \tag{S84}$$

But $d(\{x_A\})$ has the opposite sign as $q_A(\{x_A\}, T)$ (as if $x_A(T) > x_A(0)$, then $q_A > 0$) and thus the last requirement is also satisfied. All that is left for us to demonstrate convergence is to show that $V$ is positive. We do so by showing that $\nabla^2 V < 0$. This requires that

$$\frac{\partial q_A(\{x_A\}, T)}{\partial x_A^0} < 0 \tag{S85}$$

$$\iff \frac{\partial q_A(\{x_A\}, T)}{\partial x_A} \frac{\partial x_A}{\partial x_A^0} < 0 \tag{S86}$$

But $\frac{\partial x_A}{\partial x_A^0} > 0$ if $\frac{\partial q_A(\{x_A\}, T)}{\partial x_A} < 0$ so it suffices to show that

$$\frac{\partial q_A(\{x_A\}, T)}{\partial x_A} < 0 \tag{S87}$$

$$\iff 0 > \int_{s(0)}^{s(t)} \frac{\partial}{\partial x_A} \frac{r_A(s) - \bar{r}(x, s)}{ds/dt} ds \tag{S88}$$

$$\iff 0 > \int_{s(t)}^{s(0)} \frac{\partial}{\partial x_A} \frac{r_A - \bar{r}}{\sum r_\alpha \rho_\alpha} ds \tag{S89}$$

$$\iff 0 > \int_{s(t)}^{s(0)} \frac{-\bar{r}' \cdot (\sum r_\alpha \rho_\alpha) - (r_A - \bar{r})(\sum r_\alpha \rho_\alpha)'}{(\sum r_\alpha \rho_\alpha)^2} ds \tag{S90}$$

$$\iff 0 > \int_{s(t)}^{s(0)} \frac{-\bar{r}' \cdot (r_A \cdot x + \bar{r} \cdot (1 - x)) - (r_A - \bar{r})(r_A \cdot x + \bar{r} \cdot (1 - x))'}{(\sum r_\alpha \rho_\alpha)^2 / (\sum \rho_\alpha)} ds \tag{S91}$$

$$\iff 0 > \int_{s(t)}^{s(0)} \frac{\bar{r}' \cdot (r_A \cdot x + \bar{r} \cdot (1 - x)) + (r_A - \bar{r})(r_A + \bar{r}' \cdot (1 - x) - \bar{r})}{(\sum r_\alpha \rho_\alpha)^2 / (\sum \rho_\alpha)} ds \tag{S92}$$

For the case of two species, $\bar{r}' = r_B' = 0$. Thus, the condition is satisfied.

##### S2.5.3 Unique Fixed Point for Discretized Growth Functions

Assume that the range of $\rho_{\text{tot}}$ can be discretized into $N$ niches such that each species $\alpha$ (out of $M$ total species) has a constant growth rate $r_{\alpha,n}$ in niche $n < N$. If there is a stable steady state, then

$$\sum_n^N r_{\alpha,n} \tau_n = -\log \delta, \ \forall \alpha. \tag{S93}$$

where $\tau_n$ is the time spent by the community in niche $n$ in the steady cycle. Further, for each niche $n$, we have the following constraint:

$$\eta_n = \sum_{\alpha} \rho^*_{\alpha}(0) \exp(\sum_{i<n} r_{\alpha,i}\tau_i)(\exp(r_{\alpha,n}\tau_n) - 1), \tag{S94}$$

where $\rho^*_{\alpha}(0)$ is the steady cycle population density of species $\alpha$ at the beginning of the cycle. Eq. **S93** and Eq. **S94** thus give us $N + M$ constraints for the $N + M$ unknown variables ($tau_n$ and $\rho^*_{\alpha}(0)$). Thus, unless there is a strong degeneracy such that the transcendental equation in Eq. **S94** yields multiple possible solutions (for example, if $r_{\alpha,n}$ is the same for all $\alpha$ in each niche $n$), there can only be one solution to the system of equations Eq. **S93** and Eq. **S94**.

##### S2.5.4 Existence of non-trivial solutions can be inferred by mutual invasibility

Inferring when a possible frequency exists such that $q = 0$ can be very complicated because the time dependence of the growth rates cannot be inferred without solving the ODEs numerically. However, because we know there is a negative frequency dependence, we can infer if such an $x^*$ exists by asking what the values of $q(0)$ and $q(1)$ are. If $q > 0$ for all $x$, that would mean that Species A always has a fitness advantage over Species B. Similarly, if $q < 0$ for all $x$, that would mean that Species B always has a fitness advantage over Species A.

The ability to understand the necessary and sufficient conditions for coexistence by considering only the cases that either species is present (as $x = 0$ is the case when only Species B is present, and $x = 0$ is the case when only Species A is present) is very valuable when studying systems with non-linear growth rates. The study of these cases, also known as invasion analysis, is ordinarily sufficient to demonstrate coexistence. But because of the result that $q' < 0$, it is also necessary.

For our case, mutual invasibility can be verified without any numerical simulations. This is because the condition $q = 0$ can be written as $\mathbb{E}(r_A/r_B) > 1$, where $\mathbb{E}(\cdot)$ is the time-averaged expectation value. Mathematically, $\mathbb{E}(\cdot)$ is the integral of the argument from the lowest biomass value to the highest biomass value attained in the monoculture, weighted by $\omega(s) \equiv [-\log\delta \cdot (\frac{n_0}{1-\delta} - s)]^{-1}$. $\omega(s)$ is the equivalent of the partition function for biomass accumulation.

Similarly, $q(1) < 0$ can we written as $\mathbb{E}(r_B/r_A) > 1$. If both monocultures are invadable, then there necessarily is coexistence.[1]

This criterion provides a general intuition of when there is coexistence: when both $r_A/r_B$ and $r_B/r_A$ are large for different parts of the growth step, both monocultures are invadable and there is coexistence. Thus, we need a range of biomass values when A grows much faster than B, and a range when B grows much faster than A. Various trade-offs can facilitate such a separation of biomass ranges, as does the strength of the non-linearity. This indicates to us why coexistence is increased when biologically-motivated modifications are introduced.

This criterion also indicates when interactions between two species can facilitate coexistence: when the interactions between two species lead to the initially slower-growing species eventually growing faster than the initially faster-growing species. This is highlighted in Amarnath et al. [27] showing how stress-induced cross-feeding leads to stable coexistence in a marine co-culture. Species A grows faster than Species B initially, but then pollutes its environment by secreting acetate as a by-product of growth. This pollution leads to the supression of its own growth and the leakage of metabolites. Species B can thus consume these metabolites to grow. Thus, this leads to the creation of periods when A grows much faster than B, and a period when B grows much faster than A. Though the experiment was performed in growth-dilution cycles, in natural settings the periodic supply of food describes a very similar dynamic in the system. The experimental result was surprising because if the coexistence were due to the standard picture of commensal/mutualistic cross-feeding where one species steadily secretes a metabolite consumed by the other, there would be no coexistence for this pair of species as under ideal conditions, one species always grows faster than the other.

Thus, this mutual invasibility criterion provides a counter-point to the standard conception of coexistence due to mutualism in which both species promote each others' growth. In time varying environments, both species can end up limiting their own growth rates and thus coexist. This suggests that self-organized mechanisms by which species inhibit their own growth may facilitate coexistence.

[1]This also allows us to see that there cannot be bistability, as that would require that $\mathbb{E}(r_A/r_B) < 1$ and $\mathbb{E}(r_B/r_A) < 1$ but $\mathbb{E}(r_A/r_B) + \mathbb{E}(r_B/r_A) > 2$ for all $r_B$ and $r_A$ as $x + 1/x > 2$ for all $x > 0$.

##### S2.6 Mutual invasibility criterion

Here, we derive the criterion presented in Eq. **3**. Let's consider a monoculture of Species A with a minimal amount of Species B such that $(\rho_A^{(j)}(0), \rho_B(0)) = (\rho_A^0, \epsilon)$ and $\rho_A^0 \gg \epsilon$. Thus,

$$\log \frac{\rho_B^{(j+1)}(0)}{\rho_B^{(j)}(0)} = \int_0^T r_B(\rho_{\text{tot}}(t)) \cdot dt + \log \delta \tag{S95}$$

Since $\rho_{\text{tot}}$ is monotonic in $t$, we can substitute $t$ with $\rho_{\text{tot}}$

$$= \int_0^T \frac{r_B(\rho_{\text{tot}}) \cdot d\rho_{\text{tot}}}{d\rho_{\text{tot}}/dt} + \log \delta \tag{S96}$$

$$= \int_{\rho_A^0+\epsilon}^{\rho_{\text{tot}}^{\text{max}}} \frac{r_B(\rho_{\text{tot}})}{r_A(\rho_{\text{tot}}) \cdot \rho_A^{(j)}(t)(\rho_{\text{tot}}) + r_B(\rho_{\text{tot}}) \cdot \rho_B^{(j)}(t)(\rho_{\text{tot}})} d\rho_{\text{tot}} + \log \delta \tag{S97}$$

Assuming $r_A(\rho_{\text{tot}}) > 0$, we can always choose $\epsilon$ such that $\rho_B^{(j)}(\rho_{\text{tot}}) \ll \rho_A^{(j)}(\rho_{\text{tot}})$, $\forall \rho_{\text{tot}}$, and thus $\rho_B^{(j)}(\rho_{\text{tot}}) \ll \rho_A^{(j)}(\rho_{\text{tot}}) \cdot r_A(\rho_{\text{tot}})/r_B(\rho_{\text{tot}})$ and $\rho_{\text{tot}} = \rho_A^{(j)}(\rho_{\text{tot}})$

$$\approx \int_{\rho_A^0}^{\rho_{\text{tot}}^{\text{max}}} \frac{r_B(\rho_{\text{tot}})}{r_A(\rho_{\text{tot}})} \frac{d\rho_{\text{tot}}}{\rho_{\text{tot}}} + \log \delta. \tag{S98}$$

Similarly,

$$\log \frac{\rho_A^{(j+1)}(0)}{\rho_A^{(j)}(0)} \approx \int_{\rho_A^0}^{\rho_{\text{tot}}^{\text{max}}} \frac{r_A(\rho_{\text{tot}})}{r_A(\rho_{\text{tot}})} \frac{d\rho_{\text{tot}}}{\rho_{\text{tot}}} + \log \delta = \log \rho_{\text{tot}}^{\text{max}} - \log \rho_A^0 + \log \delta \tag{S99}$$

$$\implies \rho_A^{(j+1)}(0) = \delta \rho_{\text{tot}}^{\text{max}}. \tag{S100}$$

Thus, in subsequent cycles,

$$\log \frac{\rho_B^{(j+k+1)}(0)}{\rho_B^{(j+k)}(0)} \approx \int_{\delta\rho_{\text{tot}}^{\text{max}}}^{\rho_{\text{tot}}^{\text{max}}} \frac{r_B(\rho_{\text{tot}})}{r_A(\rho_{\text{tot}})} \frac{d\rho_{\text{tot}}}{\rho_{\text{tot}}} + \log \delta. \tag{S101}$$

$$\implies \log \frac{\rho_B^{(j+k+1)}(0)}{\rho_B^{(j)}(0)} \approx k \int_{\delta\rho_{\text{tot}}^{\text{max}}}^{\rho_{\text{tot}}^{\text{max}}} \frac{r_B(\rho_{\text{tot}})}{r_A(\rho_{\text{tot}})} \frac{d\rho_{\text{tot}}}{\rho_{\text{tot}}} + \int_{\rho_A^0}^{\rho_{\text{tot}}^{\text{max}}} \frac{r_B(\rho_{\text{tot}})}{r_A(\rho_{\text{tot}})} \frac{d\rho_{\text{tot}}}{\rho_{\text{tot}}} + (k+1) \log \delta \tag{S102}$$

For $k \to \infty$,

$$\rho_B^{(j+k+1)}(0) \gg \epsilon \text{ if } I_{A,B} \equiv \int_{\delta\rho_{\text{tot}}^{\text{max}}}^{\rho_{\text{tot}}^{\text{max}}} \frac{r_B(\rho_{\text{tot}})}{r_A(\rho_{\text{tot}})} \frac{d\rho_{\text{tot}}}{\rho_{\text{tot}}} > \log \delta. \tag{S103}$$

This is the invasibility criterion in Eq. **3**. If both $I_{A,B} > \log \delta$ and $I_{B,A} > \log \delta$, then there must be a non-trivial fixed point of the system, and by the proof in Section S2.5, it must be stable and further, the only stable solution.

##### S2.7 Continuous Growth Relations

###### S2.7.1 Monod Relation

A popular choice in microbiology and ecology for $r_i$ is the Monod relation [28], also known as the Michaelis-Menten function or the Holling's type II functional response. It is chosen to describe growth that is proportional to the nutrient availability at low nutrient concentrations but saturates at high nutrient concentrations to a maximal rate.

We first consider the effect of varying the two environmental parameters, $s_0$ and $\delta$ for Species A and B with growth rates as described in Fig. S4A. We simulated six-hours long growth dilution cycles. In general, we find that the system is in the vicinity of a steady-state in less than $\sim 100$ cycles (our results did not change significantly for longer cycles or for more cycles). The steady-state cycle is defined as having population densities and nutrient concentrations that are exactly the same in consecutive cycles.

As can be seen in Fig. S4A for our choice of physiological parameters ($r_i^{\max}$ and $K_i$), Species A (shown in red) has a higher growth rate than Species B (shown in blue) when the nutrient concentration is high (because $r_A^{\max} > r_B^{\max}$), while Species B has a higher growth rate when the nutrient concentration is low (because $K_A^{\max} > K_B^{\max}$). Such a trade-off is known as the opportunist-gleaner trade-off [29]. Although the empirical existence of such a trade-off is debated [30–32], we note that in this simplest case which has no other interactions, such a trade-off is necessary for coexistence as otherwise one species will always have a lower growth rate than the other and thus eventually be out-competed.

In Fig. S4B, we report the average fitness (average difference in growth rate over one cycle) of Species A over Species B after 100 cycles for different environmental parameters. A positive fitness value (denoted by red shading) indicates that the population of Species A is driving the population of Species B down, and thus Species B will eventually be removed from the system. A negative fitness value (denoted by blue shading) indicates that the population of Species A is being driven down and will eventually be removed from the system, while a near-zero fitness value (denoted by white shading) means that both species have reached a non-zero steady state population and thus will coexist indefinitely. As can be seen and as would be expected, high nutrient supply favors the species with the higher value of $r_i^{\max}$, while low nutrient supply favors the species with the lower $K_i$. Similarly, a lower dilution factor favors the species with the higher value of $r_i^{\max}$ as the relative amount of time spent in higher nutrient concentrations is higher.

We also report the average fitness of Species A over Species B for different physiological parameters and fixed environmental parameters ($\delta = 0.1, s_0 = 10K_B$) in Fig. S4C. As would be expected, if there is no opportunist-gleaner trade-off, as in the top left and bottom right quadrants of the phase plot, there would be no coexistence. Further, even if there is a trade-off, coexistence is not guaranteed as can be seen in the other two quadrants which have red, white, and blue regions. Thus, coexistence is not a trivial consequence of the trade-off. However, there is a narrow parameter regime of coexistence between the region where Species A dominates and where Species B dominates.

In 1972, Stewart and Levin showed mathematically that two species with an opportunist-gleaner trade-off may coexist indefinitely in growth-dilution cycles (Fig. 1C).They also demonstrated that such a coexistence was "structurally stable", i.e., it was robust to noise in the environmental/experimental parameters (Fig. S4B). As can be seen, this coexistence is also robust to small physiological perturbations (Fig. S4C) and there exists an entire region in the environmental and physiological parameter phase space (shaded in white) where the two species coexist, rather than just being a boundary between the two regions. This is a key result as it shows that this kind of coexistence is not a result of narrow fine-tuning, but seems to incorporate stabilizing mechanisms that promote coexistence. While this was a crucial novel result, it has been mostly dismissed in literature [29], due to the relatively small region of coexistence in parameter phase space. However, as we will see below, this region of coexistence is significantly broadened when biologically-realistic effects are considered. As such, it indicates a more general emergent principle for self-stabilized coexistence rather than serving as just a niche special case.

###### S2.7.2 Cut-off due to Maintenance Energy

Bacterial species require a certain minimum amount of energy, known as maintenance energy, to be able to grow. If sufficient non-zero amount of nutrients to generate this energy are not supplied, the bacteria cease to grow. This effect can be incorporated by subtracting a constant value of maintenance energy consumption rate, given by $J_i$, from the growth rate of each species and setting the minimal growth rate to be 0 (i.e, we exclude death). This leads to the following form for the growth rate:

$$r_i(s) = r_i^{\max} \cdot \frac{s}{s + K_i} - J_i. \tag{S104}$$

This expands the coexistence region of phase space considerably. In fact, for almost any two species defined by the physiological parameters $r_i$, $K_i$, and $J_i$, the coexistence region of the environmental phase-space occupies an entire half-plane (see Fig. S4E). This is because at very low resource concentrations, one of the species necessarily grows, while the other doesn't. The species that grows at very low resource concentrations will subsequently always survive for all environmental parameters.

The coexistence region in physiological parameter space is also extended, with species benefiting from even relatively minor differences in resource affinities. This is because strict coexistence only requires sufficient fold change between cycles rather than any minimal abundance.

##### S2.7.3 Hysteretic Growth Kinetics

We note that the Monod relation is obtained empirically for a perfectly-adapted bacterial population, i.e., the bacterial population is grown at a static nutrient concentration and its growth rate is measured for that nutrient concentration. In a growth-dilution cycle, the bacterial population may not have adapted to the nutrient concentration it experiences at any instant as the nutrient concentration is constantly changing. This can be viewed as a hysteresis in the growth-rate due to the slow adaptation of the internal state of the cells that constitute the population. Similar hysteretic effects are modeled by the Droop model which is popular in studying algae [33] and is also seen in macroscopic organisms as a predator's searching, attacking, or handling efficiency often empirically increases as prey density increases. This is because the feeding response of organisms often display some form of learning behavior, as a predator must have a minimal encounters with its prey before the predator is maximally efficient at feeding on that prey item.

Such a growth dependence is known as a Type III functional response and is described by the Hill function Eq. **S105**,

$$r_i(s) = r_i^{\max} \cdot \frac{s^k}{s^k + (K_i)^k}, \tag{S105}$$

where $k > 1$ is a positive number that denotes the number of minimal encounters the consumer must have with its food to increase its efficiency [34].

We note that the Monod relation is a special case of the Hill function (when $k = 1$), and increasing $k$ leads to an increase in the coexistence region of phase-space (see the case of $k = 2$ shown in Fig. S4D-F).

##### S2.7.4 Lag Time

Another effect that can be incorporated is the presence of a lag-time, such that both species do not grow for a short period of time at the beginning of each cycle. By design, the presence of lag times benefits the species with the shorter lag-time. This does not significantly increase (or decrease) the coexistence region of the physiological and environmental phase spaces but rather shifts it in favor of one species. We note that though previous studies [35] found coexistence and bistability for species with constant growth rates and lag times in growth-dilution cycles, these studies require the dilution step to be a bottleneck step such that the total initial population is fixed (which requires changing the dilution amount every cycle). These studies also required very large differences in the yield (of the factor of $\sim$100-1000) between the two species. In the absence of these two effects, the coexistence disappears for constant growth rates [36] but persists for nonlinear growth rates.

A result that has been indicated numerically by previous studies for the case of Monod growth [37–39] and is implicit in the phase diagrams of Fig. S4 is that there is no initial condition dependence in this system. Thus, there are only three steady-state possibilities that are determined by environmental and physiological parameters alone and hold for all initial conditions: that Species A eventually takes over the system, that Species B eventually does so, or that both species coexist indefinitely in a defined steady-state cycle. In other words, there is no bistability or multistability. We also note from Fig. S4 that coexistence is determined by all six parameters that effectively describe the system and that each parameter can be varied to lead to any of the three possible outcomes. This implies that the system cannot be described as being a trivial outcome of a simple characteristic of the environment or physiology of the two species, but requires an interplay of the environment and the physiologies. Thus, though it is necessary that the growth rate dependences of the two species intersect, it is far from sufficient.

We note that if all $r_i$ are linear in the concentration of the nutrient (also known as a Holling's type I functional response), coexistence is not possible and only one species can survive [16]. This is because one species will always grow faster than the other species and thus out-compete the other species over many cycles. Thus, nonlinear growth dependences on nutrient concentrations are required for coexistence and reflect a trade-off in the growth process.

Thus, we note that even for two species competing for a single nutrient, the coexistence region of parameter phase space is not necessarily small, but possibly a result of multiple physiological trade-offs between growth rate, resource affinity, adaptation time scale, and survivability in low resource conditions. For other systems shown in Fig. 1, the trade-offs could be due to susceptibility to a pollutant, cross-feeding, or anomalous response to environmental stress.

Accordingly, growth-dilution cycles result in a consumer-resource analog to winnerless competition models, where transient dynamics enable multiple species to persist [40].

##### S2.8 Derivation of Equi-abundance Solution

We take each species, $\alpha$, to have growth rate $r_-$ basally and $r_+$ in its preferred growth rate. In the steady cycle, we look for the solution where there are $N$ species, each with abundance $1/N$. First, since this is the steady cycle, we require

$$\sum_i r_{\alpha,i} \cdot \tau_i = \log(1/\delta), \tag{S106}$$

where $\tau_i$ is the time spent passing through each niche $i$. Plugging in the growth rates, we have

$$r_+\tau_\alpha + r_-(T - \tau_+) = \log(1/\delta), \tag{S107}$$

where $\tau_\alpha$ is the time spent in the preferred niche. Since the rest of the equation carries no parameters unique to species $\alpha$, $\tau_\alpha$ must be the same for all species. Thus, we have that

$$\tau \equiv \tau_\alpha = \frac{\log(1/\delta)}{r_+ + r_-(N-1)},\ \forall\alpha \tag{S108}$$

Now, we seek to find $\Delta\eta_n$ for each niche. Since this is the biomass consumed by all species in that niche, we have

$$\Delta\eta_n = \underbrace{\sum_{i<n} \frac{1}{N}\exp((n-2)r_-\tau + r_+\tau)(\exp(r_-\tau)-1)}_{\text{Species that have a preferred niche before the nth niche}} + \underbrace{\frac{1}{N}\exp((n-1)r_-\tau)(\exp(r_+\tau)-1)}_{\text{Species that prefers the nth niche}} \tag{S109}$$

$$+ \underbrace{\sum_{i>n} \frac{1}{N}\exp((n-1)r_-\tau)(\exp(r_-\tau)-1)}_{\text{Species that have a preferred niche after the nth niche}} \tag{S110}$$

After some simplification,

$$\Delta\eta_n = \frac{\exp((n-1)r_-\tau)(\exp(r_-\tau)-1)}{N}\left(\sum_{i<n}\exp((r_+-r_-)\tau) + \frac{\exp(r_+\tau)-1}{\exp(r_-\tau)-1} + \sum_{i>n} 1\right) \tag{S111}$$

$$\implies \Delta\eta_n = \frac{\exp((n-1)r_-\tau)(\exp(r_-\tau)-1)}{N}\left((n-1)\exp((r_+-r_-)\tau) + \frac{\exp(r_+\tau)-1}{\exp(r_-\tau)-1} + N - n\right) \tag{S112}$$

In the case that the correction term is independent of $n$, i.e.,

$$n(\exp((r_+-r_-)\tau)-1) \ll \frac{\exp(r_+\tau)-1}{\exp(r_-\tau)-1} + N - \exp((r_+-r_-)\tau), \tag{S113}$$

we have that

$$\Delta\eta_n \propto \exp(nr_-\tau) \implies \Delta\eta_n \sim \delta^{\frac{n}{p+N-1}} \tag{S114}$$

and the niche widths must be exponentially spaced. We now explore the cases where this is true. This is obviously the case if $r_+ = r_-$ as then $\exp((r_+ - r_-)\tau) - 1 = 0$.
In the case that $N \gg p \equiv \frac{r_+}{r_-} > 1$, we have that

$$\exp(r\tau) = \exp(\frac{r/r_- \cdot \log(1/\delta)}{p+N-1}) \approx 1 + \frac{r/r_- \log(1/\delta)}{N} \tag{S115}$$

Thus, the LHS in correction term is given by

$$n\left(\frac{p-1-\log(1/\delta)}{N}\right) \ll N \tag{S116}$$

while the RHS is given by

$$\frac{\frac{p-\log(1/\delta)}{N}}{\frac{1-\log(1/\delta)}{N}} + N - \frac{p-1-\log(1/\delta)}{N} - 1 \approx N. \tag{S117}$$

Thus, Eq. **S113** is satisfied. Also, in the case that $p \gg N > 1$,

$$\exp((r_+ - r_-)\tau \to \exp(r_+\tau) = \exp\left(\frac{\log(1/\delta)}{1+\frac{N-1}{p}}\right) \approx \frac{1}{\delta}. \tag{S118}$$

and

$$\exp(r_- \cdot \tau \to \exp\left(\frac{\log(1/\delta)}{p+N-1}\right) \approx 1 + \frac{\log(1/\delta)}{p+N-1}. \tag{S119}$$

Thus,

$$\Delta\eta_n = \frac{\exp((n-1)r_-\tau)}{N}\frac{\log(1/\delta)}{p+N-1}\left(n(1/\delta-1) + \frac{\frac{1}{\delta}-1}{\frac{\log(1/\delta)}{p+N-1}} + N - \frac{1}{\delta}\right) \tag{S120}$$

$$= \frac{\exp((n-1)r_-\tau)}{N}\log(1/\delta)\left(\frac{n(1/\delta-1)}{p+N-1} + \frac{\frac{1}{\delta}-1}{\log(1/\delta)} + \frac{N-\frac{1}{\delta}}{p+N-1}\right) \tag{S121}$$

$$\approx \frac{\exp((n-1)r_-\tau)}{N}\left(\frac{1}{\delta}-1\right) \propto \delta^{\frac{n}{p+N-1}} \tag{S122}$$

##### S2.9 The diagonal preference model

Based on the equiabundance solution, we consider a "diagonal model" in growth preference, in which each niche $n$ has a dominant species $\hat{\alpha}(n)$ whose growth rate $r_{\hat{\alpha}(n),n}$ well exceeds the growth rate of other species in that niche, and each species is dominant only in one unique niche.

We anticipate the equi-abundance case to be highly stable. We validated this expectation by creating ensembles of $r_{\alpha,n}$ and $\Delta\eta_n$ values fluctuating within $\pm 30\%$. Notably, every species persists in $\sim 80\%$ of scenarios with near equi-abundance (Fig. S6B, S6C). We also found that higher growth preference $p$ species in its main niche (with the accompanying change in the exponential niche distribution) increased survival rates of over 95%. In fact, even with just a 3-fold growth preference and $\pm 30\%$ noise, all species remained in 45-65% of cases. This nonlinear reliance on growth preference indicates diminishing returns.

Yet, as Fig. S6D illustrates, the proportion of preserved species for a set growth preference ($p$) declines as species and niche counts rise. The drop depends on $N/p$ (Fig. S6E), emphasizing enhanced competition from the growing collective of slow-growing species.

Alternatively, the priority effect can be overcome by having the species preferred in earlier niches take on reduced growth preferences but equal niche widths. We find that exponentially-distributed growth rates for species in their preferred niches,

$$r_+(n) \propto (1/\delta)^{n/(N+p-1)}, \tag{S123}$$

with $r_-$ fixed, retains a similar diversity (Fig. S6G, S6H) as for exponentially distributed niche widths (Fig. S6B, S6C). See Fig. S6.

Collectively, these results underscore the tremendous (exponential) growth advantage of species specializing in early niches if $p$ is not too large, and hence the much higher relative dominance required for species specializing in the late niches to be maintained in the community.

##### S2.10 Resource Sharing in Consumer-Resource Models in a Chemostat

One might interpret the states in the Community State (CS) Model as analogous to resources in a Consumer-Resource model. In the case of large communities, it leads to the question of how many species can be expected to coexist if $N$ species shared $N$ resources, with each species growing on multiple resources.

In the context of the CS Model, species coexist by occupying distinct niches, with niche width and overlap playing critical roles in determining community structure. Translating this to the CR model framework, we postulated that the allocation and consumption of resources could be a surrogate for these niche dynamics. Thus, we simulated the CR model in a chemostat. Each species was described by a consumption/growth matrix with diagonal elements set to 1 (such that each species invariably consumed a specific resource) and off-diagonal elements set randomly as 1 or 0 with the constraints described below. The growth rates, We considered two distinct cases:

$K_s = M$ Case: In this case, we fixed the number of species that show rapid growth per resource. This setup parallels a situation in the CS Model where each niche supports a similar number of species. The results can be seen in Supplementary Figure S7.

$K_n = M$ Case: In this case, we explored a scenario where each species is allocated a fixed number of resources, akin to each species in the CS Model occupying $M$ niches. The results can be seen in Supplementary Figure S8.
